# Grad-seq shines light on unrecognized RNA and protein complexes in the model bacterium *Escherichia coli*

**DOI:** 10.1101/2020.06.29.177014

**Authors:** Jens Hör, Silvia Di Giorgio, Milan Gerovac, Elisa Venturini, Konrad U. Förstner, Jörg Vogel

**Affiliations:** Institute of Molecular Infection Biology, University of Würzburg, D-97080 Würzburg, Germany; ZB MED - Information Centre for Life Sciences, D-50931 Cologne, Germany; TH Köln, Faculty of Information Science and Communication Studies, D-50678 Cologne, Germany; Helmholtz Institute for RNA-based Infection Research (HIRI), D-97080 Würzburg, Germany

**Keywords:** Grad-seq, RNA, RBP, complexome, *E. coli*

## Abstract

Stable protein complexes, including those formed with RNA, are major building blocks of every living cell. *Escherichia coli* has been the leading bacterial organism with respect to global protein-protein networks. Yet, there has been no global census of RNA/protein complexes in this model species of microbiology. Here, we performed Grad-seq to establish an RNA/protein complexome, reconstructing sedimentation profiles in a glycerol gradient for ~85% of all *E. coli* transcripts and ~49% of the proteins. These include the majority of small noncoding RNAs (sRNAs) detectable in this bacterium as well as the general sRNA-binding proteins, CsrA, Hfq and ProQ. In presenting use cases for utilization of these RNA and protein maps, we show that a stable association of RyeG with 30S ribosomes gives this seemingly noncoding RNA of prophage origin away as an mRNA of a toxic small protein. Similarly, we show that the broadly conserved uncharacterized protein YggL is a 50S subunit factor in assembled 70S ribosomes. Overall, this study crucially extends our knowledge about the cellular interactome of the primary model bacterium *E. coli* through providing global RNA/protein complexome information and should facilitate functional discovery in this and related species.

## INTRODUCTION

RNA-protein interactions leading to the formation of stable cellular complexes underlie many processes that are conserved in all domains of life. In bacteria, these traditionally range from small complexes such as the ~85 kDa signal recognition particle (SRP) to the ~450 kDa RNA polymerase (RNAP) and the giant 70S ribosome, which is a ~2.5 MDa assembly of three ribosomal RNA species and more than 50 ribosomal proteins (1). CRISPR/Cas systems involved in genome defense are a recent addition to this list and keep surprising with unexpected molecular diversity (2). In contrast with eukaryotic biology where there have been large-scale initiatives to systematically map RNA-protein interactions and complexes in different model species, it is fair to say that we have an incomplete knowledge of the breadth of macromolecular complexes in bacteria, even in primary model species such as *Escherichia coli*.

*E. coli* K-12 has been the workhorse not only for bacterial genetics and physiology but also for the identification and functional characterization of complexes of bacterial proteins, with or without RNA. There is a large body of functional in-depth studies in *E. coli* employing low-throughput biochemical assays, results of which are organized and accessible via curated databases such as EcoCyc (3). In addition, *E. coli* has been subject to large-scale affinity purification followed by mass spectrometry (AP/MS) in order to determine candidate cellular networks of protein-protein interactions (4–7). Similarly, yeast two-hybrid screens have predicted binary interactions for nearly all possible combinations within the *E. coli* proteome (8). Beyond these binary methods, the *E. coli* complexome has been studied with global methods directly, i.e., without epitope-tagging or heterologous expression of proteins (9–12). This includes recent protein-correlation-profiling, whereby complexes of membrane proteins were predicted following the “guilt-by-association” logic after their separation by size exclusion chromatography (13).

An important information that has been missing is which *E. coli* transcripts and proteins engage in stable cellular complexes, both determined in the same experiment. Such predictions can now be made in a genome-wide manner by combined high-throughput RNA-seq and mass spectrometry (MS) analyses of a bacterial lysate after its fractionation in a glycerol gradient. Referred to as gradient profiling by sequencing (Grad-seq) (14), this method predicts RNA/protein complexes by looking for co-migration of a given RNA and protein in the same gradient fraction (15). In its pioneering application (14), Grad-seq uncovered in *Salmonella enterica* the ProQ/FinO-domain protein ProQ as a previously overlooked global RNA-binding protein (RBP) that impacts the expression of several hundred different transcripts (16–18).

Here, we use Grad-seq to provide an RNA/protein complexome resource for *E. coli*. By reporting sedimentation profiles for the vast majority of cellular transcripts and half of the *E. coli* proteins, we faithfully reproduce hundreds of known protein-protein and RNA-protein interactions including major ribonucleoprotein particles (RNPs). We describe use cases for this resource to be exploited: a stable association with the 30S ribosomal subunit reveals the seemingly noncoding RyeG RNA of prophage origin as an mRNA that encodes a small toxic protein; the broadly conserved uncharacterized protein YggL is revealed to be a 50S subunit factor in assembled 70S ribosomes. The *E. coli* Grad-seq data are viewable in an online browser (https://helmholtz-hiri.de/en/datasets/gradseqec/) that not only enables easy access to all recorded RNA and protein gradient sedimentation profiles but also permits cross-species comparison with published *Salmonella enterica* (14) and *Streptococcus pneumoniae* (19) datasets.

## MATERIALS AND METHODS

### Bacteria and media

For all experiments, *Escherichia coli* MG1655 was streaked on LB plates and grown overnight at 37°C. Overnight cultures of single colonies were grown in 2 ml LB at 37°C with shaking at 220 rpm. The next day, 1:100 dilutions in fresh LB of the overnight cultures were used to start the cultures for the experiments and grown to an OD_600_ of 2.0 (early stationary phase) at 37°C with shaking at 220 rpm. All strains and plasmids are listed in Supplementary Tables S3 and S4.

### Strain construction

Gene inactivation mostly followed a published protocol (20). Briefly, a strain carrying the pKD46 plasmid, which carries the λRED recombinase and is temperature-sensitive, was grown overnight at 28°C. The next day, the overnight culture was diluted 1:300 in 50 ml LB containing 0.2% L-arabinose and grown at 28°C to an OD_600_ of 0.5. Electrocompetent cells were prepared and transformed with 800-1,000 ng of a gel-purified PCR product containing a kanamycin resistance cassette. The PCR product was obtained using the pKD4 plasmid and primers containing the flanking regions of the gene of interest. The transformed cells were streaked on LB agar plates and incubated at 37°C overnight. The deletion was verified by PCR. 3xFLAG-tagging of genes followed a published protocol (21), which is similar to the one described for gene inactivation above, except that the PCR product was obtained using the pSUB11 plasmid. Removal of the antibiotics resistance cassettes was performed by transformation of the temperature-sensitive pCP20 plasmid that carries the Flp recombinase (20). All oligos are listed in Supplementary Table S5.

### Glycerol gradient fractionation

Glycerol gradient fractionation was performed as previously described (19), with the exception of the growth and lysis conditions: 100 ml of *E. coli* MG1655 wild type were grown to an OD_600_ of 2, cooled down in an ice-water bath for 15 min and then harvested by centrifugation for 20 min at 4°C and 4,000 rcf. The cells were washed three times in ice-cold 1x TBS, resuspended in 500 μl ice-cold 1x lysis buffer A [20 mM Tris-HCl, pH 7.5, 150 mM KCl, 1 mM MgCl_2_, 1 mM DTT, 1 mM PMSF, 0.2% Triton X 100, 20 U/ml DNase I (Thermo Fisher), 200 U/ml RNase inhibitor] and lysed by addition of 750 μl of 0.1 mm glass beads (Carl Roth) and 10 cycles of vortexing for 30 s followed by cooling on ice for 15 s. To remove insoluble debris and the glass beads, the lysate was cleared by centrifugation for 10 min at 4°C and 16,100 rcf.

Of the cleared lysate, 10 μl was mixed with 1 ml TRIzol (Thermo Fisher) for the RNA input control and 20 μl was mixed with 20 μl 5x protein loading buffer for the protein input control. 200 μl of the cleared lysate was then layered on top of a linear 10-40% (w/v) glycerol gradient (in 1x lysis buffer A without DNase I or RNase inhibitor), which was formed in an open-top polyallomer ultracentrifugation tube (Seton Scientific) using the Gradient Station model 153 (Biocomp). The gradient was centrifuged for 17 h at 4°C and 100,000 rcf (23,700 rpm) using an SW 40 Ti rotor (Beckman Coulter), followed by manual fractionation into 20 590 μl fractions and measurement of the A_260 nm_ of each fraction. 90 μl of each fraction and 40 μl of the pellet were mixed with 30 μl of 5x protein loading buffer for protein analysis and stored at −20°C.

The remaining 500 μl of each fraction were used for RNA isolation by addition of 50 μl of 10% SDS (25 μl for the pellet) and 600 μl of acidic phenol/chloroform/isoamylalcohol (P/C/I; 300 μl for the pellet). The fractions were then vortexed for 30 s and let rest at room temperature for 5 min before separating the phases by centrifugation for 15 min at 4°C and 16,100 rcf. The aqueous phases were collected, 1 μl of GlycoBlue (Thermo Fisher) and 1.4 ml of ice-cold ethanol/3 M sodium acetate, pH 6.5 (30:1) were added and precipitated for at least 1 h at −20°C. The RNA was collected by centrifugation for 30 min at 4°C and 16,100 rcf and washed with 350 μl ice-cold 70% ethanol, followed by centrifugation for 15 min at 4°C and 16,100 rcf. The lysate RNA sample stored in TRIzol was purified according to the manufacturer’s protocol, except that the precipitation was performed using the mentioned ethanol mix. After drying of the RNA pellet, it was dissolved in 40 μl DEPC-treated H_2_O and DNase-digested by addition of 5 μl DNase I buffer with MgCl_2_ (Thermo Fisher), 0.5 μl RNase inhibitor, 4 μl DNase I and 0.5 μl DEPC-treated H_2_O, followed by incubation for 45 min at 37°C. The DNase-treated RNA was purified by addition of 150 μl DEPC-treated H_2_O and 200 μl P/C/I as described above. The purified, DNase-treated RNA was dissolved in 35 μl DEPC-treated H_2_O and stored at −80°C.

### Sucrose polysome gradient fractionation

50 ml of *E. coli* MG1655 was grown to an OD_600_ of 2, followed by rapid filtration and immediate freezing in liquid N_2_. The cells were then resuspended in 1 ml of ice-cold 1x lysis buffer B [20 mM Tris-HCl, pH 7.5, 100 mM NH_4_Cl, 10 mM MgCl_2_, 1 mM DTT, 1 mM PMSF, 0.4% Triton X 100, 20 U/ml DNase I, 200 U/ml RNase-inhibitor] and lysed using a FastPrep-24 instrument (MP Biomedicals) and a 2 ml lysing matrix E tube (MP Biomedicals) for 15 s at 4 m/s. To remove insoluble debris and the beads, the lysate was cleared by centrifugation for 10 min at 4°C and 16,100 rcf. Of the cleared lysate, 10 μl was mixed with 1 ml TRIzol for the RNA input control. 15 A_260 nm_ per ml of the cleared lysate were then layered on top of a linear 10-55% (w/v) sucrose gradient (in 1x lysis buffer B without DNase I or RNase inhibitor and with addition of 5 mM CaCl_2_), which was formed in an open-top polyclear ultracentrifugation tube (Seton Scientific) using the Gradient Station model 153. The gradient was centrifuged for 2.5 h at 4°C and 237,000 rcf (35,000 rpm) using an SW 40 Ti rotor, followed by automated fractionation into 20 fractions using an FC 203B fractionator (Gilson). RNA extraction was performed as for the glycerol gradient, except that the vortexing step was performed for 15 s and that DNase treatment of the purified RNA was skipped.

### RNA gel electrophoresis and northern blotting

Equal volumes of the gradient RNA samples (glycerol or sucrose) were separated by 6% denaturing PAGE in 1x TBE and 7 M urea and stained with ethidium bromide. For northern blotting, unstained gels were transferred to Hybond+ membranes (GE Healthcare Life Sciences) and probed with RNA-specific radioactively labeled DNA oligonucleotides.

### Protein gel electrophoresis and western blotting

Equal volumes of the gradient protein samples were separated by 12% SDS-PAGE and stained with Coomassie. For western blotting, unstained gels were transferred to PVDF membranes (GE Healthcare Life Sciences) and probed with protein-specific antisera against 3xFLAG (Sigma-Aldrich, cat# F1804), 6xHis (Sigma-Aldrich, cat# H1029), GroEL (Sigma-Aldrich, cat# G6532), RpoB (BioLegend, cat# 663905) or RpoD (BioLegend, cat# 663202). Visualization of the primary antibodies was performed using anti-mouse (Thermo Fisher, cat# 31430) or anti-rabbit (Thermo Fisher, cat# 31460) secondary antibodies.

### RNA-seq

RNA-seq was performed as described before (19). Briefly, 5 μl of the gradient samples were diluted in 45 μl DEPC-treated H_2_O. 10 μl of the resulting 1:10 dilution were mixed with 10 μl of a 1:100 dilution of the ERCC spike-in mix 2 (Thermo Fisher) and subjected to library preparation for next-generation sequencing (vertis Biotechnologie). Briefly, the RNA samples were fragmented using ultrasound (4 pulses of 30 s at 4°C) followed by 3’ adapter ligation. Using the 3’ adapter as primer, first strand cDNA synthesis was performed using M-MLV reverse transcriptase. After purification, the 5’ Illumina TruSeq sequencing adapter was ligated to the 3’ end of the antisense cDNA. The resulting cDNA was PCR-amplified to about 10-20 ng/μl using a high-fidelity DNA polymerase followed by purification using the Agencourt AMPure XP kit (Beckman Coulter). The cDNA samples were pooled with ratios according to the RNA concentrations of the input samples and a size range of 200-550 bp was eluted from a preparative agarose gel. This size-selected cDNA pool was finally subjected to sequencing on an Illumina NextSeq 500 system using 75 nt single-end read length.

### RNA-seq data analysis

Pre-processing steps like read trimming and clipping were done with cutadapt (22). Read filtering, read mapping, nucleotide‐wise coverage calculation, and genome feature‐wise read quantification was done using READemption (23) (v0.4.3; https://doi.org/10.5281/zenodo.250598) and the short read mapper segemehl (24) (v0.2.0‐418) with the *Escherichia coli* MG1655 genome (accession number NC_000913.3) as reference. The annotation provided was extended by ncRNAs predicted by ANNOgesic (25). The analysis was performed with the tool GRADitude (Di Giorgio, S., Hör, J., Vogel, J., Förstner, K.U., unpublished; v0.1.0; https://foerstner-lab.github.io/GRADitude/). Only transcripts with a sum of ≥ 100 reads in all fractions within the gradient were considered for the downstream analysis. Read counts for each fraction, were normalized by calculating size factors following the DESeq2 approach (26) generated from the ERCC spike‐in read counts added to each sample (see above). To remove left‐over disturbances in the data, the size factors were then manually adjusted by multiplication based on quantified northern blots: 1.5 (fraction 5), 4.5 (fraction 7) and 28 (fraction 8). In order to make all the transcript counts comparable, they were scaled to the maximum value.

After normalization, analyses based on the detected transcripts were performed. t‐SNE dimension reduction (27) was performed using the Python package scikit‐learn (28). All default parameters provided by the sklearn.manifold.TSNE class were used. A file collection representing the analysis workflow, including Unix Shell calls, Python scripts, documentation as well as resulting files have been deposited at Zenodo (https://doi.org/10.5281/zenodo.3876866).

### Sample preparation for mass spectrometry

Sample preparation for mass spectrometry was performed as described before (19). Briefly, the gradient protein samples (diluted in 1.25x protein loading buffer) were homogenized using ultrasound [5 cycles of 30 s on followed by 30 s off, high power at 4°C (Bioruptor Plus, Diagenode)]. Insoluble material was then removed by centrifugation for 15 min at 4°C and 16,100 rcf. 20 μl of the cleared protein sample were mixed with 10 μl of UPS2 spike-in (Sigma-Aldrich) diluted in 250 μl 1.25x protein loading buffer. The samples were then reduced in 50 mM DTT for 10 min at 70°C and alkylated with 120 mM iodoacetamide for 20 min at room temperature in the dark. The proteins were precipitated in four volumes of acetone overnight at −20°C. Pellets were washed four times with acetone at −20°C and dissolved in 50 μl 8 M urea, 100 mM ammonium bicarbonate. Digestion of the proteins was performed by addition of 0.25 μg Lys-C (Wako) for 2 h at 30°C, followed by dilution to 2 M urea by addition of 150 μl 100 mM ammonium bicarbonate and overnight digestion with 0.25 μg trypsin at 37°C. Peptides were desalted using C-18 Stage Tips (29). Each Stage Tip was prepared with three disks of C-18 Empore SPE Disks (3M) in a 200 μl pipet tip. Peptides were eluted with 60% acetonitrile/0.3% formic acid, dried in a laboratory freeze-dryer (Christ) and stored at −20°C. Prior to nanoLC-MS/MS, the peptides were dissolved in 2% acetonitrile/0.1% formic acid.

### NanoLC-MS/MS analysis

NanoLC-MS/MS analysis was performed as described before (19) using an Orbitrap Fusion (Thermo Scientific) equipped with a PicoView Ion Source (New Objective) and coupled to an EASY-nLC 1000 (Thermo Scientific). Peptides were loaded on capillary columns (PicoFrit, 30 cm x 150 μm ID, New Objective) self-packed with ReproSil-Pur 120 C18-AQ, 1.9 μm (Dr. Maisch) and separated with a 140 min linear gradient from 3% to 40% acetonitrile and 0.1% formic acid at a flow rate of 500 nl/min. Both MS and MS/MS scans were acquired in the Orbitrap analyzer with a resolution of 60,000 for MS scans and 15,000 for MS/MS scans. HCD fragmentation with 35% normalized collision energy was applied. A Top Speed data-dependent MS/MS method with a fixed cycle time of 3 s was used. Dynamic exclusion was applied with a repeat count of 1 and an exclusion duration of 60 s; singly charged precursors were excluded from selection. Minimum signal threshold for precursor selection was set to 50,000. Predictive AGC was used with a target value of 2×10^5^ for MS scans and 5×10^4^ for MS/MS scans. EASY-IC was used for internal calibration.

### MS data analysis

MS data analysis was performed as described before (19), with a few exceptions. Raw MS data files were analyzed with MaxQuant version 1.5.7.4 (30). The search was performed against the UniProt database for *E. coli* MG1655 (organism identifier: ECOLI), a database containing the UPS2 spike-in and a database containing common contaminants. The search was performed with tryptic cleavage specificity with 3 allowed miscleavages. Protein identification was under control of a false-discovery rate of 1% on both protein and peptide level. In addition to the MaxQuant default settings, the search was performed against the following variable modifications: Protein N-terminal acetylation, Gln to pyro-Glu formation (N-terminal Q) and oxidation on Met. For protein quantitation, the LFQ intensities were used (31). Proteins with less than 2 identified razor/unique peptides were dismissed.

Normalization of the proteins across the fractions was performed using the UPS2 spike-in. For this, only spike-in proteins with detectable intensities in all fractions were used. The spike-in proteins showing the highest variance (median average deviation of log10 intensities >1.5x lQR) were eliminated. Following this, for each spike-in protein, the median log10 intensity was subtracted from the log10 intensities of each fraction. The fraction-wise median of the resulting values was then subtracted from the log10 intensities for each bacterial protein in the corresponding fractions. Finally, all log_10_ intensities smaller than the 5% quantile of all intensities in the data set were replaced by the value of the 5% quantile of all intensities in the data set.

### *Estimation of* in vivo *RNA copy numbers*

Estimation of *in vivo* copy numbers of RyeG was performed as described previously (32). Briefly, RNA was extracted at the given time points by collecting 4 OD_600_ of cells. The RNA was diluted in 40 μl water and 10 μl from each time point (≈10^9^ cells) were probed on a northern blot. For reference, *in vitro*-transcribed RyeG was loaded (0.05, 0.1, 0.5, 1 and 2.5 ng). RNA levels per cell were based on determination of viable cell counts per OD_600_ as described in (33).

### 30S subunit toeprinting analysis

30S subunit toeprinting was performed as previously published (34,35) with few changes. Briefly, 0.2 pmol unlabeled, *in vitro*-transcribed RyeG and 0.5 pmol of a 5’-labeled DNA oligonucleotide (JVO-16833) were denatured for 1 min at 95°C in the presence of 0.8 μl SB 5x -Mg (50 mM Tris-acetate, pH 7.6, 500 mM potassium acetate, 5 mM DTT) in a total volume of 3 μl. After incubation on ice for 5 min, 1 μl dNTPs (5 mM each) and 1 μl SB 1x Mg60 (10 mM Tris-acetate, pH 7.6, 100 mM potassium acetate, 1 mM DTT, 60 mM magnesium acetate) were added and the samples were incubated for 5 min at 37°C. Next, 4 pmol purified 30S subunits (pre-activated for 20 min at 37°C) was added to the samples [SB 1x Mg10 (10 mM Tris-acetate, pH 7.6, 100 mM potassium acetate, 1 mM DTT, 10 mM magnesium acetate) was added to the control]. After incubation for 5 min at 37°C, 10 pmol uncharged fMet-tRNA^Met^i was added to the corresponding sample. Reactions were continued at 37°C for 15 min, followed by addition of 100 U SuperScript II reverse transcriptase (Thermo Fisher) and incubation for 20 min at 37°C.

Reactions were stopped by addition of 100 μl toeprint stop buffer (50 mM Tris-HCl, pH 7.5, 0.1% (w/v) SDS, 10 mM EDTA, pH 8). DNA was extracted by addition of 110 μl P/C/I. Next, 5 μl 3 M KOH was added and the RNA digested at 90°C for 5 min. 10 μl 3 M acetic acid, 1 μl GlyoBlue and 300 μl ethanol/3 M sodium acetate, pH 6.5 (30:1) were added and the DNA precipitated at −20°C overnight. Extraction was finished and the pellet washed once with 100 μl of 70% ethanol. The purified pellet was dissolved in 10 μl 1x RNA loading buffer, denatured for 3 min at 90°C and subjected to separation using a denaturing 8% sequencing gel in presence of a RyeG-specific sequencing ladder prepared using the DNA Cycle Sequencing kit (Jena Bioscience) according to the manufacturer’s instructions. Gels were run for 1.5 h at 40 W, dried and exposed on a phosphor screen.

### Purification of ribosomes

Crude purification of ribosomes mostly followed a previously published protocol (36). Briefly, 800 ml of an *E. coli* Δ*yggL* culture was grown to an OD_600_ of ∼0.5–0.7 and washed once with 25 ml of ice-cold 1x TBS. The cell pellets were snap-frozen in liquid nitrogen and stored at −80 °C. The pellets were then resuspended in 6 ml ice-cold lysis buffer C (20 mM Tris-HCl, pH 7.5, 100 mM NH_4_Cl, 10.5 mM MgCl_2_, 0.5 mM EDTA, 3mM DTT) on ice. Lysis was performed by two lysis steps using a french press at 10,000 psi. 75 μl of 100 mM PMSF was added and the lysates cleared by centrifugation for 30 min at 4°C and 30,000 rcf using an SW 40 Ti rotor. 12.5 ml of the supernatant was subsequently layered on top of a 12.5 ml 1.1 M sucrose cushion made up in lysis buffer C. Next, the sample was centrifuged for 16 h at 4°C and 100,000 rcf using a type 70 Ti rotor (Beckman Coulter). The pellet was gently washed with 500 μl of storage buffer (lysis buffer C + 10% (v/v) glycerol) and finally dissolved in 1 ml of storage buffer by gentle shaking for 2.5 h at 4°C. After centrifugation for 5 min at 16,100 rcf and 4°C, the concentration was measured, the purified ribosomes aliquoted, snap-frozen in liquid nitrogen and stored at −80°C.

### *Analysis of* in vitro-*reconstituted complexes*

To test binding of YggL to purified ribosomes, given amounts of recombinant YggL (produced by the Recombinant Protein Expression core unit at the Rudolf Virchow Center in Würzburg) were mixed with purified ribosomes extracted from the Δ*yggL* strain. Then, the volume was increased to 200 μl with lysis buffer C and the samples were incubated for 10 min at 30°C with shaking at 330 rpm to allow complex formation. The samples were subsequently loaded on 10-40% (w/v) sucrose gradients (in 20 mM Tris-HCl, pH 7.5, 100 mM NH_4_Cl, 10 mM MgCl_2_, 3 mM DTT), which were formed in open-top polyclear tubes. Gradients were centrifuged for 14 h at 4°C and 71,000 rcf (20,000 rpm) using an SW 40 Ti rotor. Fractionation was performed as described for the sucrose gradients above.

### Co-immunoprecipitation of YggL followed by MS

Cells representing 50 OD_600_ of the *yggL*-3xFLAG or wild-type strains were collected and washed once with 1 ml of lysis buffer A. After resuspension in 800 μl lysis buffer A, the cells were transferred to a 2 ml FastPrep tube with lysing matrix E and lysed using a FastPrep-24 instrument for 20 s at 4 m/s. The lysate was cleared for 10 min at 16,100 rcf and 4°C. 40 μl of magnetic protein A/G beads (Thermo Fisher) were washed with 1 ml of lysis buffer A, resuspended in 400 μl lysis buffer A and 3 μl anti-FLAG antibody was added. After rotating for 45 min at 4°C, the beads were washed twice with 400 μl lysis buffer A. 600 μl of the lysate was added to the beads with the coupled antibody and rotated for 1.5 h at 4°C. The beads were washed five times with 400 μl lysis buffer A and briefly spun down. The lysis buffer was removed, the beads were resuspended in 35 μl 1x LDS sample buffer (Thermo Fisher) with 50 mM DTT and the proteins eluted by incubation at 95°C for 5 min.

The samples were alkylated in presence of 120 mM iodoacetamide for 20 min in the dark and run on a precast 4-12% Bolt Bis-Tris plus gel (Thermo Fisher) using 1x MES buffer (Thermo Fisher). The gel was stained with SimplyBlue Coomassie (Thermo Fisher) and each lane of the gel was either cut into 11 pieces or specific prominent bands were cut. To prepare the gel pieces for LC/MS-MS, they were destained with 30% acetonitrile in 100 mM ammonium bicarbonate, pH 8. Next, the pieces were shrunk using 100% acetonitrile and dried. Digestion was performed by addition of 0.1 μg trypsin per gel piece and incubation overnight at 37°C in 100 mM ammonium bicarbonate, pH 8. The supernatant was removed and the peptides were extracted from the gel pieces with 5% formic acid. Finally, the supernatant was pooled with the extracted peptides and subjected to MS.

## RESULTS

### *Grad-seq reveals sedimentation of the soluble RNA and protein content of* E. coli

We performed Grad-seq on *E. coli* grown to early stationary phase (OD_600_ of 2.0) in rich medium. As before (14,19), soluble particles from a lysate were separated on a linear 10%-40% glycerol gradient, followed by RNA-seq and MS analyses of all 20 gradient fractions and the pellet (Figure 1A). The A_260_ UV profile of the gradient showed the typical three peaks, one bulk peak around low molecular weight (LMW) fraction 2 and the two peaks representing the small (30S) and large (50S) ribosomal subunits (Figure 1B) (14,19). Fully assembled 70S ribosomes sedimented in the pellet (P). RNA gel analysis showed tRNAs, 16S rRNA and the 5S/23S rRNAs to individually overlap with those three major UV peaks, as expected (Figure 1C). Other abundant house-keeping RNAs forming stable RNPs such as 6S RNA, tmRNA or RnpB (the RNA part of RNase P (37)), were also readily detected in their expected fractions, confirming RNA integrity. Likewise, SDS-PAGE of the extracted proteins confirmed intactness of several of those major RNPs, exemplified by co-sedimentation of RNA polymerase (RNAP) proteins with 6S RNA (38) and ribosomal proteins with rRNAs (Figure 1D). The position of RNAP was further refined by western blot detection of its β-subunit (RpoB) and its major σ factor, σ^70^ (RpoD) (Figure 1E).

**Figure 1.**
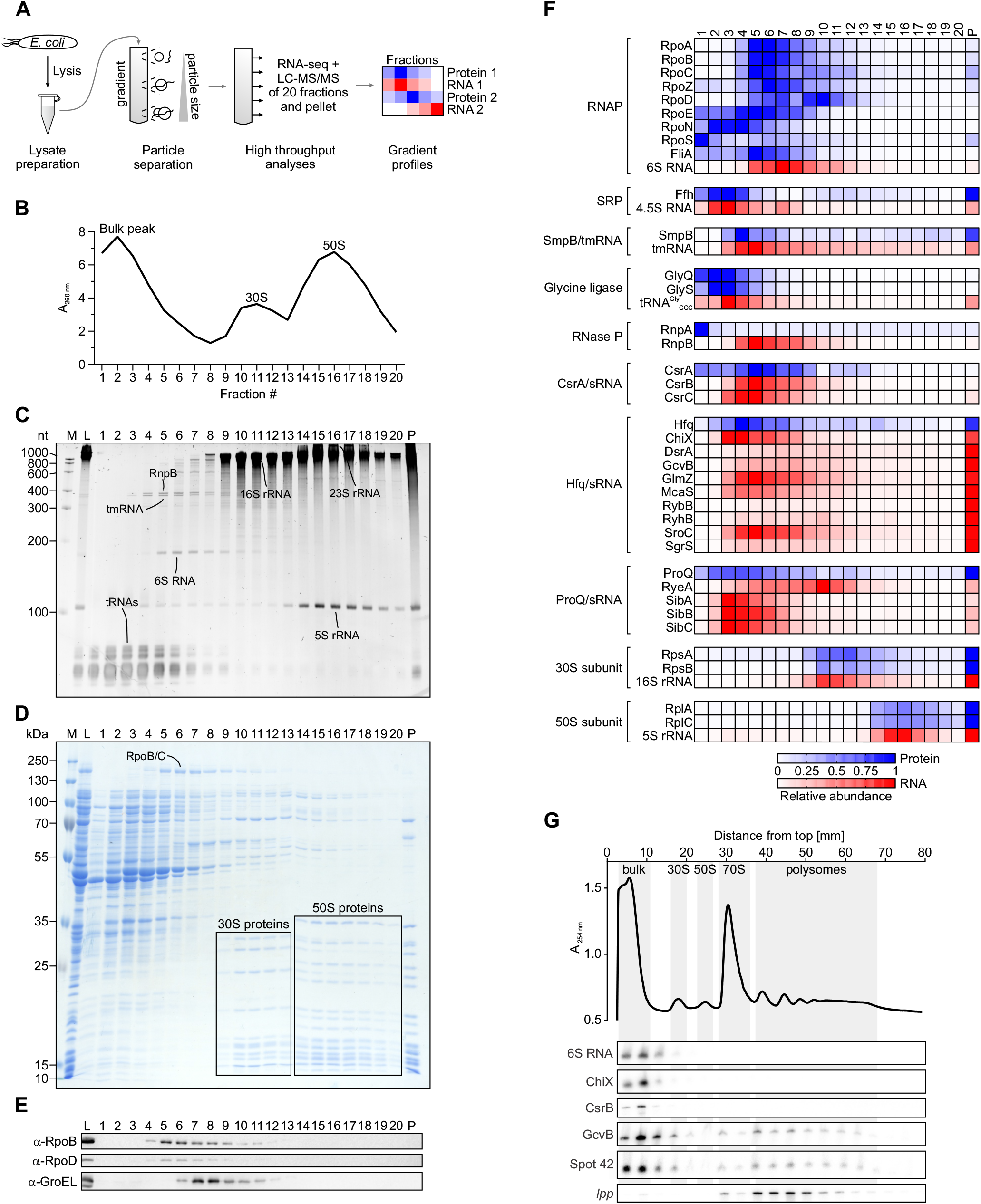
Grad-seq reveals the *E. coli* RNA/protein complexome. (A) Overview of the Grad-seq workflow. (B) A_260 nm_ profile of the gradient. Low‐molecular‐weight complexes (bulk peak) and ribosomal subunits (30S, 50S) are highlighted. Particles larger than the 50S subunit were pelleted. (C) Ethidium bromide‐stained RNA gel. Bands corresponding to abundant housekeeping RNAs are indicated. (D) Coomassie‐stained SDS–PAGE. Bands corresponding to abundant housekeeping proteins are indicated. (E) Western blot. The β-subunit of RNAP (RpoB) and the major σ factor σ^70^ (RpoD) co-migrate. (F) Heat map of digital in‐gradient distributions of known RNA–protein complexes derived from RNA‐seq and LC‐MS/MS data. The profiles are normalized to the range from 0 to 1. M, size marker. L, lysate (input control). P, pellet fraction. (G) Sucrose polysome gradient of a wild-type lysate followed by northern blotting. 6S RNA, ChiX and CsrB are only present in the bulk peak, whereas GcvB and Spot 42 show additional abundances around the polysomes (compare to (F)). The *lpp* mRNA is only present in ribosomal fractions.

### *Digital Grad-seq recovers the majority of* E. coli *transcripts and proteins*

RNA-seq of the 20 fractions and the pellet reported the sedimentation profiles of 4,095 transcripts, comprising 3,699 mRNAs, 287 ncRNAs and all tRNAs and rRNAs (Supplementary Table S1). Similar to *Salmonella* Grad-seq (14), mRNAs strongly co-sedimented with the 70S ribosome (Supplementary Figure S1A), whereas ncRNAs showed disparate behaviors (Supplementary Figure S1B). The well-known class of Hfq-binding sRNAs (39) mostly sedimented in medium-sized complexes around fraction 5, with an additional peak in the pellet (Supplementary Figure S1C). ProQ-binding sRNAs peaked earlier, around fraction 4 (Supplementary Figure S1D). In contrast, the CsrA antagonists CsrB and CsrC were confined to the gradient region around fraction 5 (Supplementary Figure S1E). Northern blotting confirmed the sedimentation profiles of the RNA-seq (average Spearman’s ≈ 0.88), thereby verifying our dataset (Supplementary Figure S2).

Parallel MS analysis detected a total of 2,145 proteins with high confidence, representing ~49% of the proteome as annotated on UniProt (40) (Supplementary Table S2). The majority of proteins sedimented in LMW fractions, suggesting involvement in no or small complexes (Supplementary Figure S1F). Of all detected proteins, ~71% have known or predicted cytoplasmic and periplasmic localization, indicating enrichment of soluble proteins during our sample preparation (Supplementary Figure S3A and B).

### Grad-seq visualizes RNPs

The possibility of analyzing complex formation of RNAs is one of the main benefits of Grad-seq compared to other methods (15). RNAP consisting of the α-, β-, β’- and ω-subunits (RpoA, RpoB, RpoC and RpoZ, respectively) co-migrated with the noncoding 6S RNA (Figure 1F), which controls transcription by competing for promoter binding of RNAP-σ^70^ (41). We note that in the MS analysis σ^70^ (RpoD) shows an intriguing second peak around fraction 10, outside RNAP, which was not detected in western blot analysis (Figure 1E). In addition, while σ^70^, σ^24^ (RpoE) and σ^28^ (FliA) generally occurred in the same fractions as did RNAP, this was only true for a fraction of the measured σ^54^ (RpoN) and σ^S^ (RpoS) intensities (Figure 1F). The ribosomal subunits, the SRP (consisting of 4.5S RNA and Ffh (42)) and the SmpB-tmRNA RNP (43) are other major RNPs for which the RNA and protein components showed excellent correlation. In contrast, RnpA being the protein factor of RNase P (37) was barely detected in the first 3 fractions, away from RnpB. This resembles previous results with *Salmonella* Grad-seq (14) and argues that RNase P has a tendency to disintegrate under the present Grad-seq conditions.

The three major regulatory RBPs of *E. coli*—CsrA, Hfq and ProQ—are known to bind specific subsets of ncRNAs (16,18,44,45). CsrA exhibited a broader peak, perhaps caused by its associations with diverse target mRNAs (46) in addition to its complexes with the major CsrB and CsrC RNAs (Figure 1F). The absence of CsrA from the pellet fraction (containing 70S ribosomes) underscores its main function as an RBP that inhibits mRNA translation.

Hfq is responsible for most sRNA-based regulation in *E. coli* (39,47,48). In our Grad-seq data, Hfq showed peaks in fraction 4 and the pellet (Figure 1F), echoing early biochemical results from even before the sRNA-related major function of Hfq was discovered (49). In general, this pattern was also seen with the Hfq-binding sRNAs (Supplementary Figure S1C). Individual sRNAs, however, exhibited disparate sedimentation profiles (Figure 1F). For example, ChiX, which is both abundant and perhaps the strongest Hfq binder (50), peaked in fraction 4 and was found in the pellet to some degree. In contrast, GcvB was almost exclusively present in the pellet fraction. This suggested that GcvB was preferentially associated with ribosomes and/or ribosome-associated Hfq. Testing this prediction, we probed for GcvB on a northern blot of a sucrose instead of a glycerol gradient, and indeed found this sRNA to be abundant in both, the 70S monosome and polysome fractions (Figure 1G).

ProQ is the least understood of the three sRNA-associated RBPs. In our *E. coli* Grad-seq data, ProQ-binding sRNAs showed a higher average abundance toward the top of the gradient around fraction 4 (Supplementary Figure S1D), which is also where ProQ was found to peak (Figure 1F). Antitoxins of type I toxin-antitoxin (TA) systems are noncoding antisense RNAs that form a well-characterized class of ProQ ligands (14,16,18) and function by repressing the translation of their corresponding toxins (51–53). Of these antitoxins, SibA, SibB and SibC coincided with ProQ, whereas RyeA, which was proposed to serve as an antitoxin to SdsR (54), sedimented away from ProQ (Figure 1F). We note that, similar to Hfq, ProQ was abundant in the pellet, supporting an earlier report of ProQ association with polysomes (55).

For a bird’s eye view of possible *in vivo* complexes of other RBPs, we filtered the MS data for proteins with predicted RNA-binding properties based on UniProt (40) and Gene Ontology (56,57) information (Supplementary Figure S4). Interestingly, these known and predicted RBPs populated the whole gradient, revealing that some are likely to act without a stable partner, whereas others are involved in complexes of different sizes. As a general observation predictive of function, we note that ribosomal proteins were generally most abundant in the pellet (where 70S ribosomes sediment), whereas proteins involved in ribosome maturation were not found in the pellet and rather co-sedimented with the 30S or 50S subunit fractions, or found elsewhere.

### RNA sedimentation profiles give functional insight

To learn more about the in-gradient behavior of RNA molecules, we performed t-stochastic neighbor embedding (t-SNE; (27)) to globally cluster all detected transcripts (Figure 2A). As expected, tRNAs as well as the rRNAs of the 30S or 50S subunits each accumulated, showing that t-SNE correctly identified their respective sedimentation profiles to be almost congruent. sRNAs mostly clustered in proximity to tRNAs, whereas mRNAs populated the whole map, as previously observed in *Salmonella* (14).

**Figure 2.**
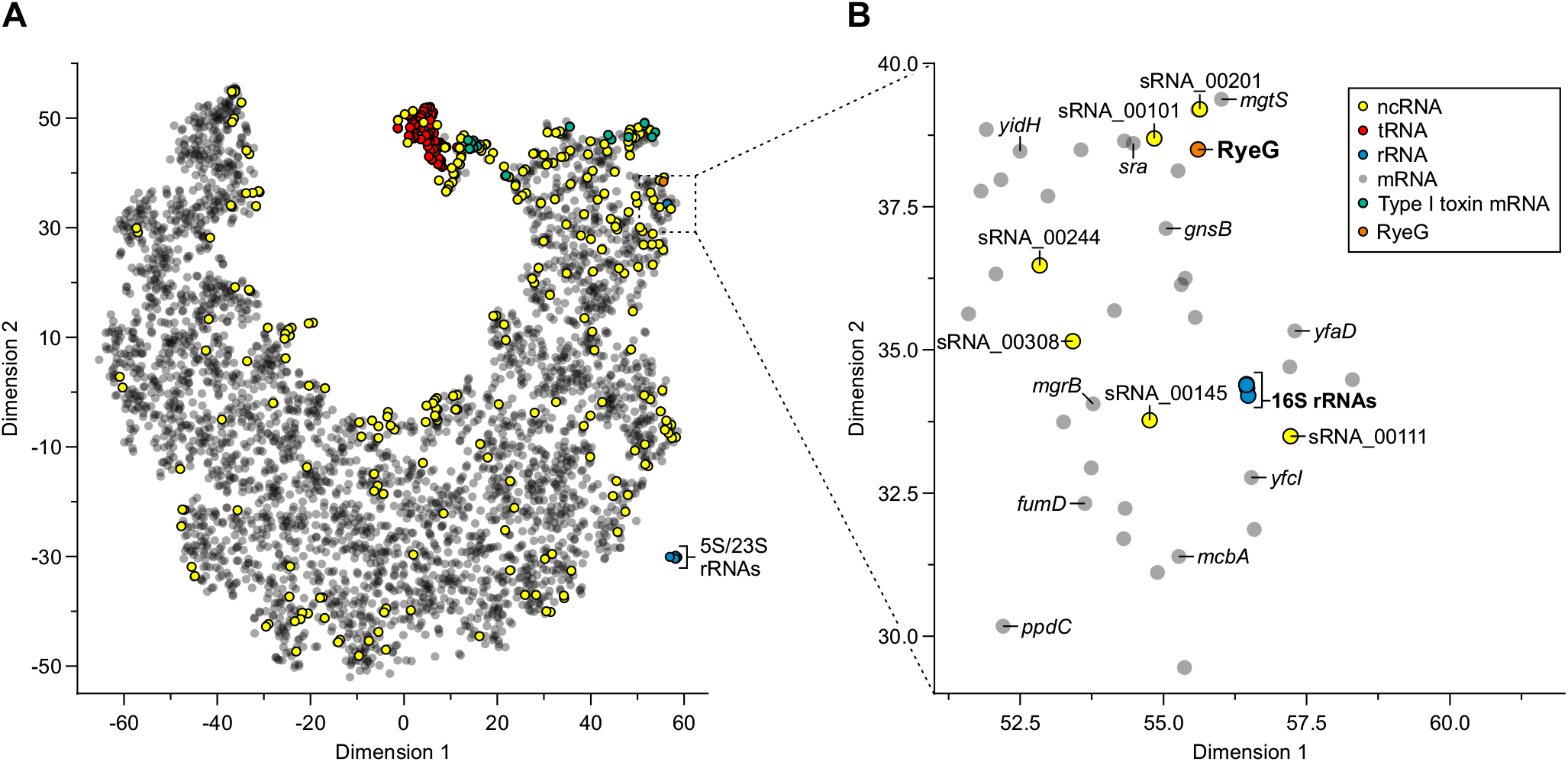
Global analysis of transcript sedimentation reveals transcripts with unexpected properties. (A) t-SNE plot of all transcripts detected in the gradient. (B) Zoomed-in region of the plot shown in (A). RyeG is in close proximity to the 16S rRNAs, indicating similar sedimentation behavior.

Interestingly, in between the 16S rRNAs and the tRNAs, we found all 16 of the toxin mRNAs of type I TA systems we detected in the gradient (Figure 2A, green). Unlike the typical mRNA (Supplementary Figure S1A), these toxin mRNAs seem to be generally excluded from 70S ribosomes, i.e., they are not found in the pellet (Supplementary Figure S5A). *Prima vista*, the prominent co-migration of antitoxin RNAs with their respective toxin mRNAs would suggest they present translationally inactive RNA-RNA complexes (Supplementary Figure S5A). However, such RNA-RNA complexes of type I TA systems are substrates of RNase III (51,52) and thus unlikely to be stable. Therefore, we interpret this co-migration to represent association with ProQ (Figure 1F) (14,16,18). In contrast to type I TA systems, both the toxins and the antitoxins of type II TA systems are proteins, meaning that the antitoxins have to be translated in order to combat the harmful effects of the toxins (53). Consequentially, both partners of type II TA systems were found to have their peak abundance in the pellet fraction (Supplementary Figure S5B).

### RyeG is a prophage-encoded RNA that co-sediments with the 30S subunit

Our t-SNE map (Figure 2A) placed several transcripts close to the 16S rRNAs, suggesting co-migration with 30S ribosomes (Figure 2B). Among these, we noticed many mRNAs coding for small proteins such as *mgrB* (58), *mgtS* (59) or *sra* (a.k.a. *rpsV*; ribosomal protein S22). Indeed, compared to all CDSs, the median length of the CDSs of these mRNAs was significantly shorter (282 aa vs. 107 aa; Supplementary Figure S5C). Furthermore, our t-SNE map revealed several ncRNAs to co-sediment with the 30S subunit. Of these, RyeG was the only sRNA that had previously been used in studies analyzing libraries of sRNA-overexpressing strains (60–63), whereas the others were low-confidence predictions. Northern blot detection of RyeG in the gradient fractions recapitulated the 30S association as well as a weaker signal in the pellet (Figure 3A-B), indicating that this noncoding RNA undergoes translation.

**Figure 3.**
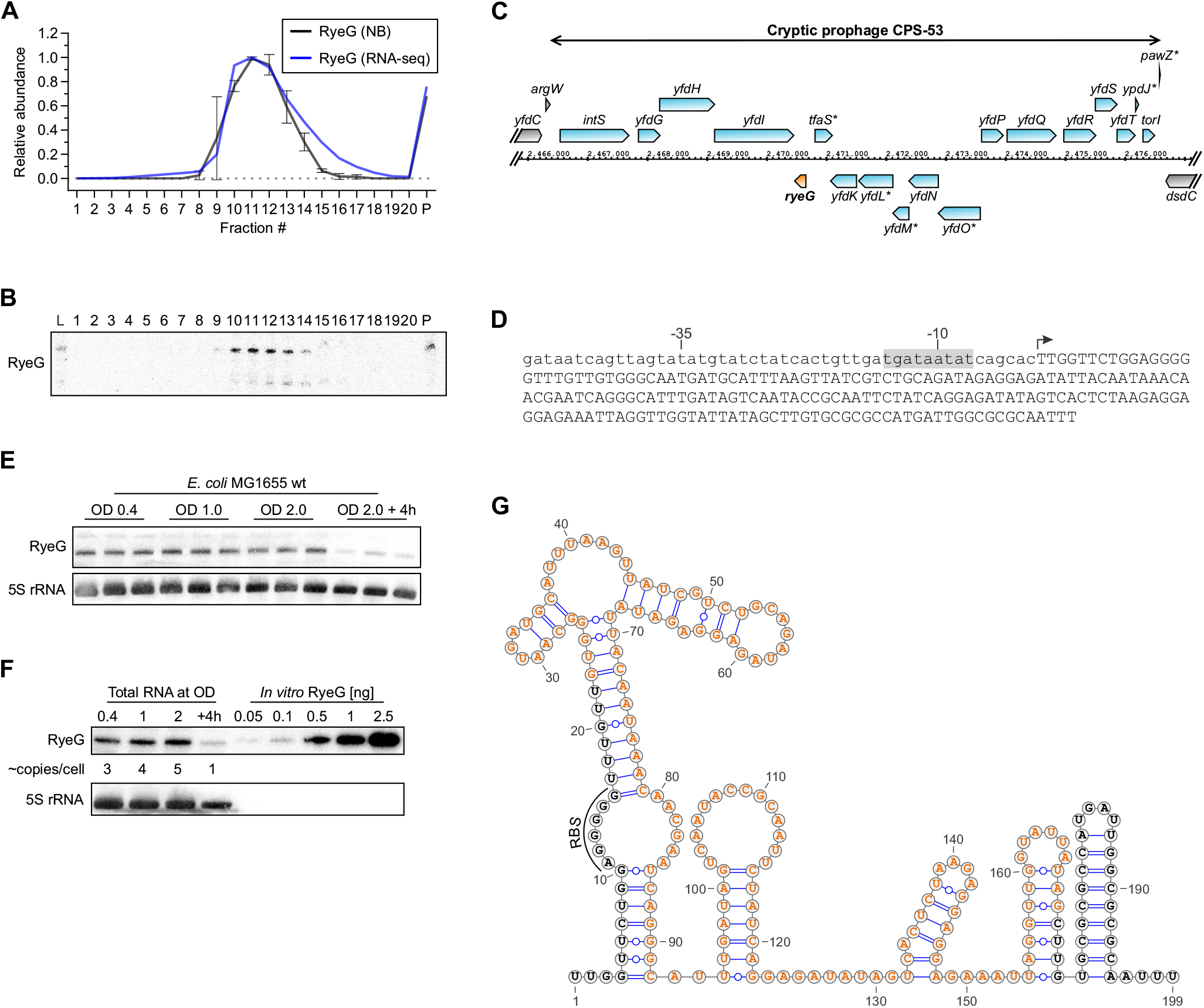
RyeG is a prophage-encoded transcript that binds the 30S subunit. (A) Sedimentation profile of RyeG. RyeG sediments around the 30S subunit. Northern blot (NB) data are quantified from (B) and a replicate. n = 2. (B) Verification of the sedimentation profile of RyeG by northern blotting. (C) Genetic locus of *ryeG*. Genes within the prophage CPS-53 are shown in blue. Genes outside of CPS-53 are shown in gray. *ryeG* is shown in orange. Asterisks denote pseudogenes. (D) Sequence of the *ryeG* locus. The predicted extended −10 box is highlighted in gray. The transcriptional start site (TSS) is indicated by an arrow. Lower case letters indicate the sequence upstream of the TSS, whereas capital letters indicate the sequence of RyeG. (E) RyeG expression during growth. Northern blotting of RyeG shows that its expression is constant during growth and is downregulated at late stationary phase. RNA from three independent biological replicates was loaded. 5S rRNA was used as loading control. (F) Estimation of *in vivo* copy numbers of RyeG. Northern blotting of RyeG compared to *in vitro*-synthesized RyeG reveals low levels of ~1-5 copies/cell. 5S rRNA was used as loading control. (G) Predicted secondary structure of RyeG. Secondary structure prediction by RNAfold (100) reveals several stem loops and a ρ-independent terminator. Nucleotides highlighted in orange represent the coding sequence of ORF2 and its ribosome-binding site (RBS) is indicated (compare to Figure 4C). Visualization was performed using VARNA (101).

RyeG was first reported as IS118 in an early bioinformatics sRNA search in *E. coli* (64) and later shown to decrease biofilm formation (62) and motility (63) when overexpressed. The *ryeG* gene lies within the cryptic prophage CPS-53, on the antisense strand between *yfdI* and *tfaS* (Figure 3C). The CPS-53 prophage is only present in *E. coli* K-12 strains and shows signs of gene erosion (65). CPS-53 has been reported to increase H_2_O_2_ and acid resistance (66) and to possess genes that inhibit initiation of chromosomal replication when overexpressed (67). The *ryeG* gene carries an extended −10 box (68) indicative of transcription by the *E. coli* housekeeping RNAP-σ^70^ (Figure 3D). Northern blot analysis showed that RyeG accumulates to 3-5 copies per *E. coli* cell during growth in rich medium, dropping to ~1 copy/cell in late stationary phase (Figure 3E-F). This 199 nt long sRNA is predicted to be highly structured (Figure 3G), which might also explain its recently described interaction with ProQ (16).

### RyeG encodes a toxic small protein

To assess whether *ryeG* was a functional gene, we overexpressed it from a high copy plasmid and determined effects on bacterial growth (Figure 4A). We observed a much longer lag-phase, with RyeG overexpressing cells reaching mid-log phase ~2 h later than wild-type *E. coli* grown in parallel. Most importantly, this growth retardation was also observed when RyeG was overexpressed in *Salmonella* (Figure 4B), which lacks CPS-53. Thus, the bacteriostatic RyeG effect is independent of any other prophage-encoded genes.

**Figure 4.**
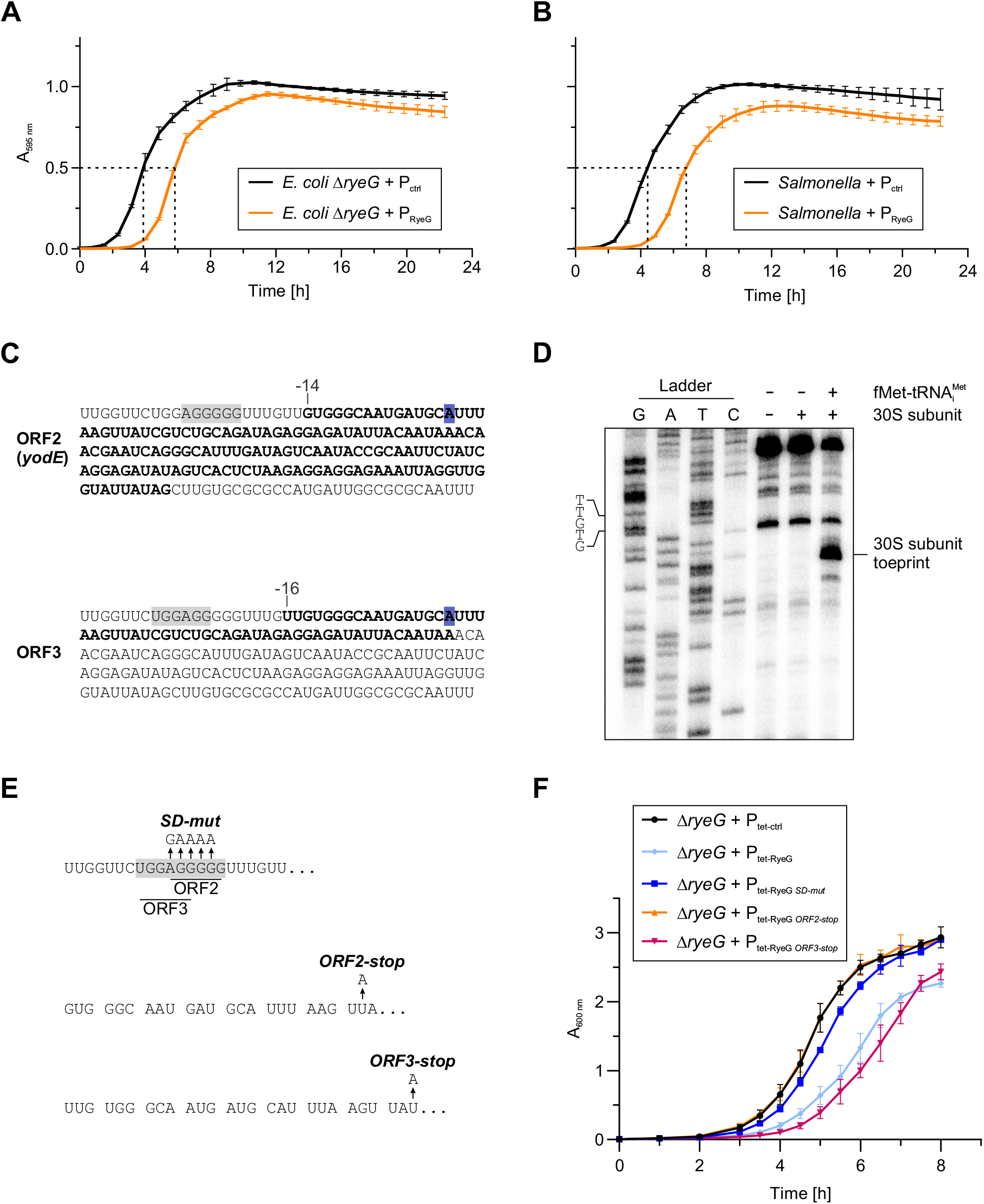
RyeG encodes for a toxic small protein. (A-B) Growth curves comparing *E. coli* Δ*ryeG* and wild-type *Salmonella* against their corresponding RyeG overexpression strains. Overexpression of RyeG leads to strongly prolonged lag times. Data was obtained from three biological replicates. (C) Sequence of RyeG with ORF2 (recently designated *yodE* (69)) and ORF3 highlighted in bold (related to Supplementary Figure S6). The corresponding predicted ribosome-binding sites are highlighted in gray. The A nucleotide at which a stop was detected using 30S subunit toeprinting (D) is highlighted in blue. (D) 30S subunit toeprinting of RyeG. A specific toeprint in presence of initiator tRNA and 30S subunits can be detected by a strong stop at an A nucleotide at position +37 of RyeG, indicating formation of an initiation complex. The overlapping start codons of ORF2 and ORF3 are indicated. (E) Mutants of RyeG. To test which ORF is responsible for the toxic effect of RyeG, three mutants were created: *SD-mut* eliminates the ribosome-binding sites of ORF2 and ORF3 without changing the predicted secondary structure (compare to Figure 3G). *ORF2-stop* and *ORF3-stop* introduce stop codons in codon 8 and 9 of ORF2 and ORF3, respectively, by single nucleotide exchanges. (F) Growth curves comparing *E. coli* Δ*ryeG* with a control plasmid against different inducible RyeG versions (E). Expression of *SD-mut* and *ORF2-stop* show similar growth as the control, whereas *ORF3-stop* shows a similar phenotype as expression of wild-type RyeG, indicating that ORF2 is responsible for the phenotype of RyeG expression. Induction of the tetracycline-inducible plasmids was performed by addition of 200 ng/ml doxycycline to the medium at the start of the experiment. Data was obtained from three biological replicates. Error bars show SD from the mean.

Next, we followed up on the observed strong 30S association of RyeG, asking whether the RNA itself or an unrecognized open reading frame (ORF) was responsible for the observed toxicity. The ORFfinder algorithm (https://www.ncbi.nlm.nih.gov/orffinder/) returned five different possible small ORFs (Supplementary Figure S6A-E) in RyeG, of which ORF2 (48 aa) and ORF3 (19 aa) were preceded by a potential ribosome-binding site (RBS) (Figure 4C). To experimentally test translation initiation, we performed toeprinting assays (34) using *in vitro*-synthesized RyeG and purified 30S subunits (Figure 4D). This revealed a strong toeprint at position +37 in presence of 30S and charged tRNA^fMet^ but not without the initiator tRNA, indicating assembly of an initiation complex. The adenosine at position +37 is located 14 nt or 16 nt upstream of ORF2 or ORF3, respectively (Figure 4C), suggesting that both ORFs can be translated, in principle.

To pinpoint the ORF that is being translated and whether it causes the observed toxicity, we constructed three different mutant versions of RyeG (Figure 4E). These had to be cloned in a tetracycline-inducible plasmid since some of them were impossible to maintain in a constitutive overexpression plasmid. As before, wild-type RyeG strongly delayed growth when expressed from the tetracycline-inducible plasmid (Figure 4F). This growth phenotype was largely abrogated by the RBS mutant (*SD-mut*), and fully so by a premature stop codon in ORF2 (*ORF2-stop*). In contrast, a premature stop codon in ORF3 (*ORF3-stop*) did not alleviate toxicity of RyeG; if at all, we observed a longer lag time than with wild-type RyeG. Therefore, RyeG is a previously unrecognized mRNA encoding a 48 aa growth-inhibitory protein. Of note, a recent global survey of candidate small proteins also showed ORF2 of RyeG to be translated, designating it *yodE* (69). This ORF2 must be exceptionally toxic, for all our attempts to clone it on its own, even under control of a tight tetracycline-dependent promoter, have failed thus far.

### Grad-seq resolves a wide range of protein complexes

Our focus on interactions of RNAs notwithstanding, Grad-seq also enables the analysis of multi-protein complexes. For many such complexes, we observed well-correlated profiles of their corresponding subunits (Figure 5A). For example, the succinyl-CoA synthetase consisting of SucC and SucD partitioned as a small complex around fraction 3, whereas the >900 kDa FtsH/HflKC metalloprotease complex sedimented as a particle almost the size of 30S subunits, matching a previous observation (70). This illustrates the wide range of complexes resolved by Grad-seq. For a more global assessment of the quality of such predictions, all heterocomplexes, for which all subunits could be detected in the MS data, were tested for co-sedimentation. Of those 107 heterocomplexes, 79 (~74%) showed high correlation (Spearman’s ≥ 0.7), indicating intact complexes (Figure 5B). Thus, following the “guilt-by-association” logic, Grad-seq profiles might be able to predict whether a given *E. coli* protein is part of a cellular complex. For orientation, a <20 kDa protein with a slightly elongated shape will sediment around <3S (71), i.e., at the top of the gradient. Conversely, if a protein <20 kDa occurs in higher fractions, it is likely involved in a complex.

**Figure 5.**
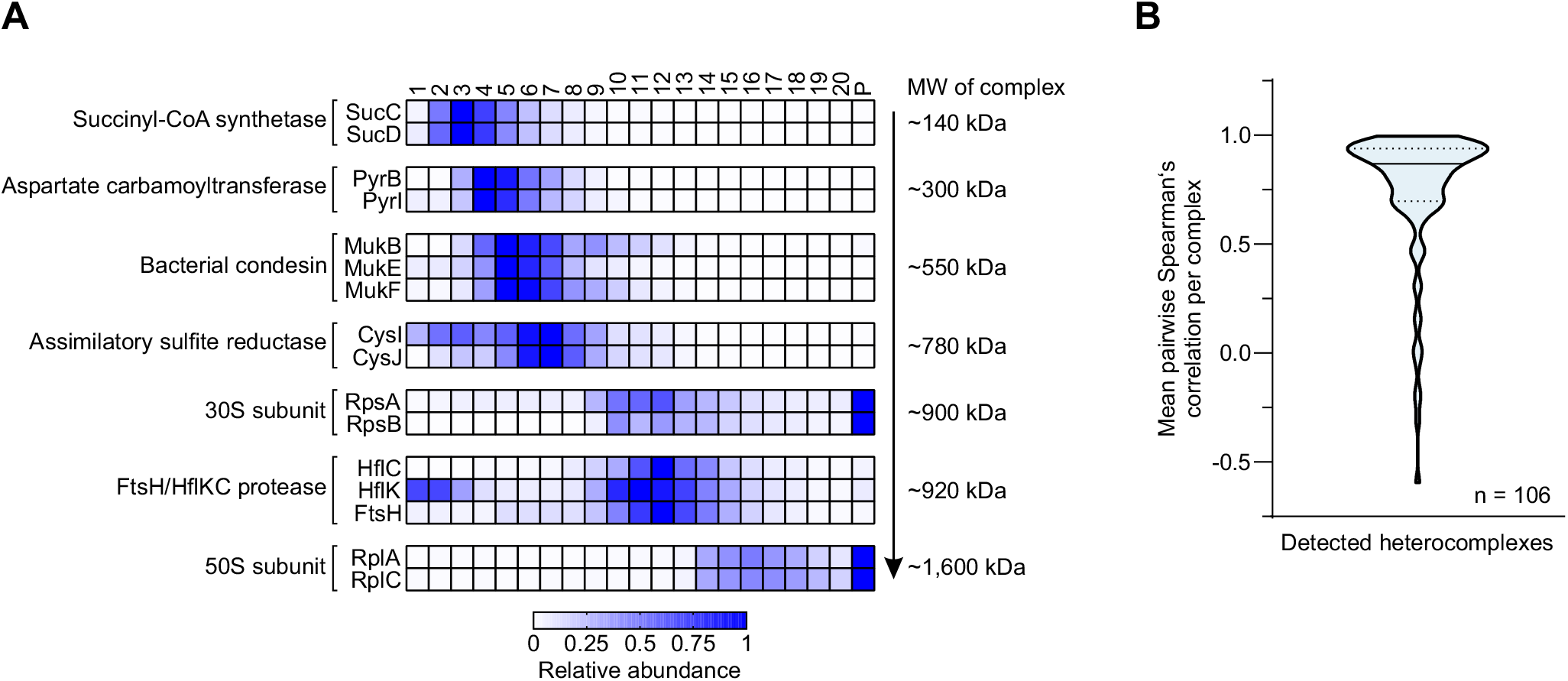
Grad-seq resolves a wide range of protein-protein complexes. (A) Heat map showing the sedimentation profiles of exemplary intact complexes spanning ~140-1,600 kDa. (B) Violin plot showing the distribution of the mean Spearman’s correlation of all heterocomplexes, for which all subunits were detected in the gradient (n = 106). 79 of these show a correlation ≥ 0.7, indicating intact complexes. The solid line indicates the median, whereas the dashed lines indicate the upper and lower quartiles.

To predict new complexes, we filtered our MS data (proteins <20 kDa with a peak in fraction ≥4) to obtain 97 proteins with unexpected in-gradient occurrences (Figure 6). Unsurprisingly, 42 of these were ribosomal proteins, and an additional four known to be ribosome-associated: Hsp15 (HslR), Rmf, RsfS and the L31 paralog YkgM. Hsp15 and RsfS co-sedimented with the 50S subunit, as reported earlier (72–75). YkgM could only be detected in fraction 15, overlapping with the height of the 50S subunit peak. Given its probable function as an alternative L31 protein (76), this may reflect a tight 50S association of YkgM. In contrast, Rmf primarily occurred in the pellet fraction, which agrees with its function in the formation of inactive 70S dimers, so-called 100S ribosomes (77,78). Other expected proteins included the RNAP-interacting proteins RpoZ, GreB (a transcription elongation factor) (79) and CedA (a regulator of cell division) (80), all of which co-sedimented with RNAP (Figure 1F). The membrane-associated proteins Lpp, OmpX and SecG were almost exclusively detected in the pellet fraction, indicating the formation of insoluble aggregates (Figure 6).

**Figure 6.**
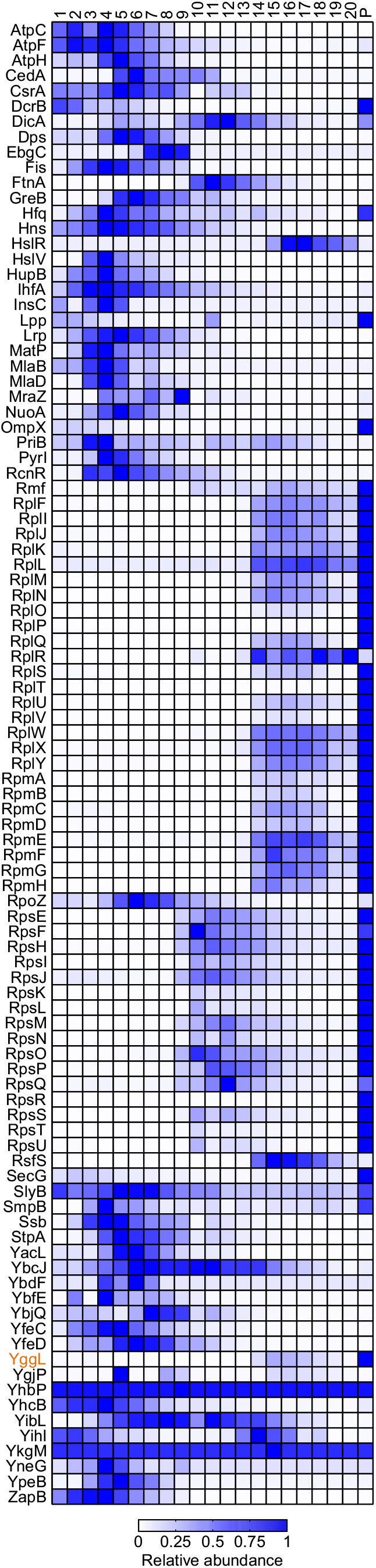
Small proteins are involved in large complexes. Heat map showing the sedimentation profiles 97 proteins <20 kDa whose peak abundance is detected in fraction 4 or higher. The profile of YggL (highlighted in orange) is congruent with the proteins of the large ribosomal subunit (Rpl* and Rpm* proteins).

### A ribosome-associated function of protein YggL

Searching for proteins with unrecognized association with larger complexes, we homed in on YggL. This small ~12 kDa protein constitutes its own family of DUF469 proteins (81), is extremely conserved in the class of γ-proteobacteria and is further present in the orders of Burkholderiales and Neisseriales within the class of β-proteobacteria (Figure 7B and Supplementary Figure S7A). According to our global MS data, YggL was most abundant in the pellet but showed an additional broad peak similar to 50S subunit components (Figure 6). To test this global MS-based prediction, we chromosomally tagged the *yggL* gene with a 3xFLAG epitope and performed western blot analysis on two different sorts of gradient samples of this *yggL*-3xFLAG strain. These analyses verified both the 50S (Figure 7B) and 70S association of YggL (Figure 7C).

**Figure 7.**
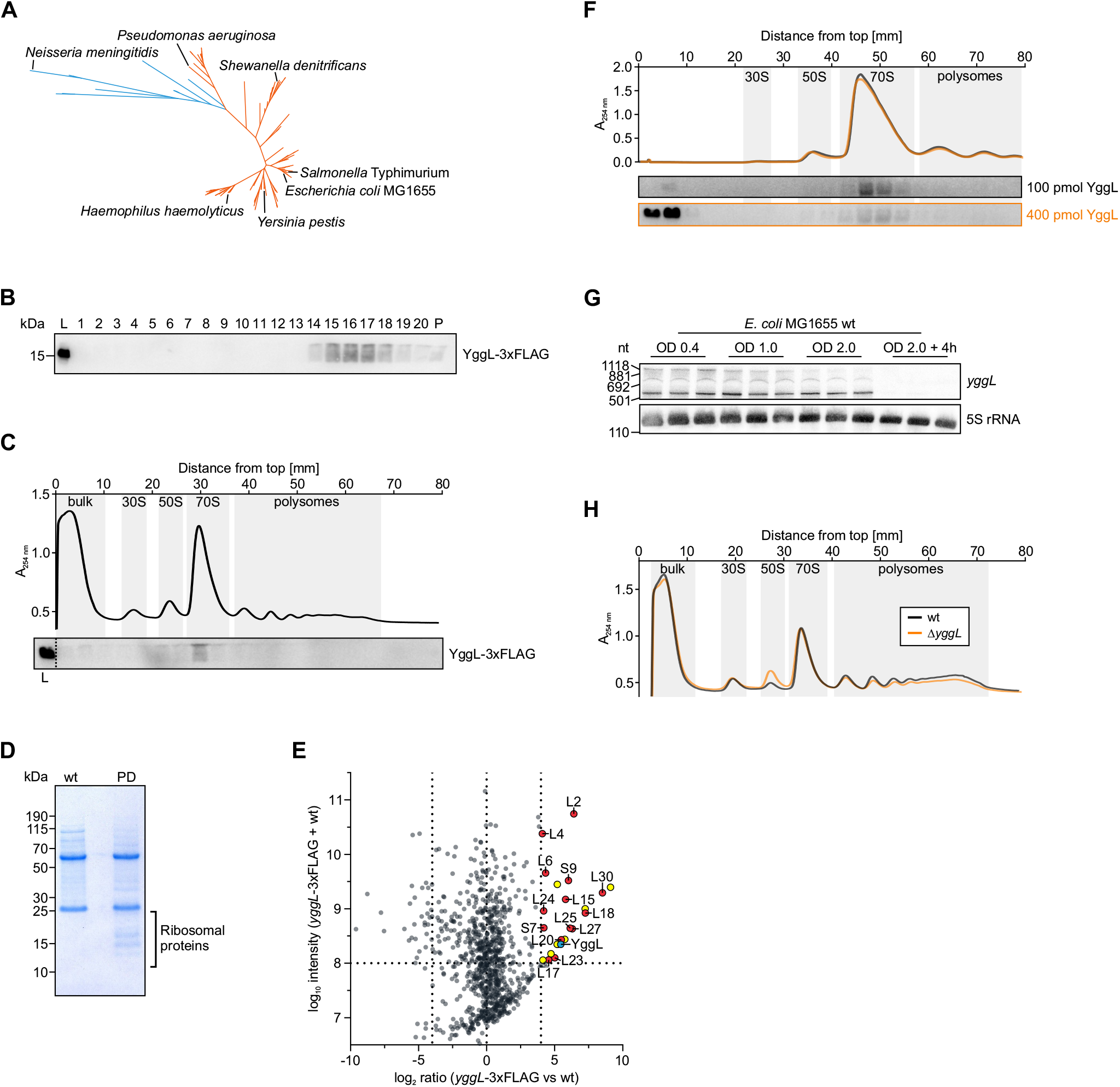
YggL is a 70S ribosome-interacting protein. (A) Phylogenetic analysis of YggL based on 150 protein sequences deposited in eggNOG 4.5.1 (COG3171; (102)). YggL homologs were only found in γ-proteobacteria (orange) and β-proteobacteria (blue). (B) Glycerol gradient analysis of a *yggL*-3xFLAG strain followed by western blotting. YggL-3xFLAG co-migrates with the 50S subunit and is found in the pellet. L, lysate (input control). P, pellet. (C) Sucrose polysome gradient analysis of a *yggL*-3xFLAG strain followed by western blotting. YggL-3xFLAG co-migrates exclusively with the 70S ribosome. L, lysate (input control). (D) SDS-PAGE of YggL-3xFLAG co-immunoprecipitation (PD) and the wild type (wt). Specific bands for the PD between 10 and 25 kDa can be detected. (E) MS analysis of the co-immunoprecipitation shown in (E). Apart from the expected enrichment of YggL (blue), several proteins of the large ribosomal subunit (red) as well as two of the small ribosomal subunit (red) were enriched compared to the wild type. Other enriched proteins are shown in yellow and proteins not considered enriched are shown in gray. (F) Sucrose polysome gradient analysis of *in vitro*-reconstituted YggL-ribosome complexes followed by western blotting. 100 pmol (black) or 400 pmol (orange) recombinant YggL was allowed to bind to 400 pmol of purified ribosomes obtained from a Δ*yggL* strain. Subsequent sucrose polysome gradient analysis shows that YggL specifically binds to 70S ribosomes. Probing for purified YggL was performed using an antibody against its 6xHis-tag, which was used for the purification of YggL and not cleaved off. (G) *yggL* expression during growth. Northern blotting of *yggL* shows that its expression is constant during growth and is shut off at late stationary phase. RNA from three independent biological replicates was loaded. 5S rRNA was used as loading control. Note that the same membrane as in Figure 3E was used, explaining the recurrence of the control lanes. (H) Sucrose polysome gradient analysis of wild-type and Δ*yggL* strains. A_260 nm_ profiles show that the knockout of *yggL* increases the amount of free 50S subunits.

The ribosome association of YggL found support in additional experiments. First, immunoprecipitation of YggL-3xFLAG from an *E. coli* lysate (Figure 7D) strongly enriched 12 proteins of the 50S subunit (Figure 7E). Interestingly, with the exception of L2 (RplB), all of them were from the ribosome’s cytosolic side (82), suggesting this is the side where YggL binds as well. Second, we confirmed the YggL-70S association by *in vitro* reconstitution of purified YggL with purified ribosomes obtained from a Δ*yggL* strain run in a sucrose gradient (Figure 7F).

Next, we followed up earlier predictions by others (83) that YggL might be involved in late 50S subunit assembly or final maturation of the 70S ribosome. This would predict YggL expression to be highest in fast-growing cells. Indeed, we observed *yggL* to be expressed only until the early stationary phase of growth (Figure 7G). Furthermore, we constructed an *E. coli* Δ*yggL* strain to test a loss-of-function effect on ribosomes. Comparing profiles of polysome gradients (Figure 7H), Δ*yggL* bacteria exhibited a strong increase in free 50S subunits, as compared to the wild-type strain. This change in cellular ribosome composition is unlikely to be a growth effect, since the *yggL* knockout grew indistinguishably from wild-type *E. coli* (Supplementary Figure S7B). Taken together, based on its Grad-seq profile, YggL emerges as a 70S ribosome-binding protein with a 50S-centric function.

### Data visualization and accessibility

Grad-seq provides a global overview of RNA and protein interactions obtained from a single experiment. To facilitate data accessibility und usability, we set up an online browser (https://helmholtz-hiri.de/en/datasets/gradseqec/) for interactive exploration of these datasets (Figure 8). Visualization can be performed based on a user-selected group of genes, displayed as bar chart, line plot or heat map and downloaded as editable vector graphics. Importantly, the browser also allows to view Grad-seq data for *Salmonella* (14) and *S. pneumoniae* (19), which permits a cross-species comparison of the sedimentation profiles of selected entries. Comparison of the closely related enterobacteria *E. coli* and *Salmonella* whose proteins tend to be generally similar will be useful for a fine-grained analysis of in-gradient distributions. Comparing potential complex formation of distantly related homologs of Gram-negative and Gram-positive bacteria may give clues for broadly conserved functions of a protein or RNA molecule.

**Figure 8.**
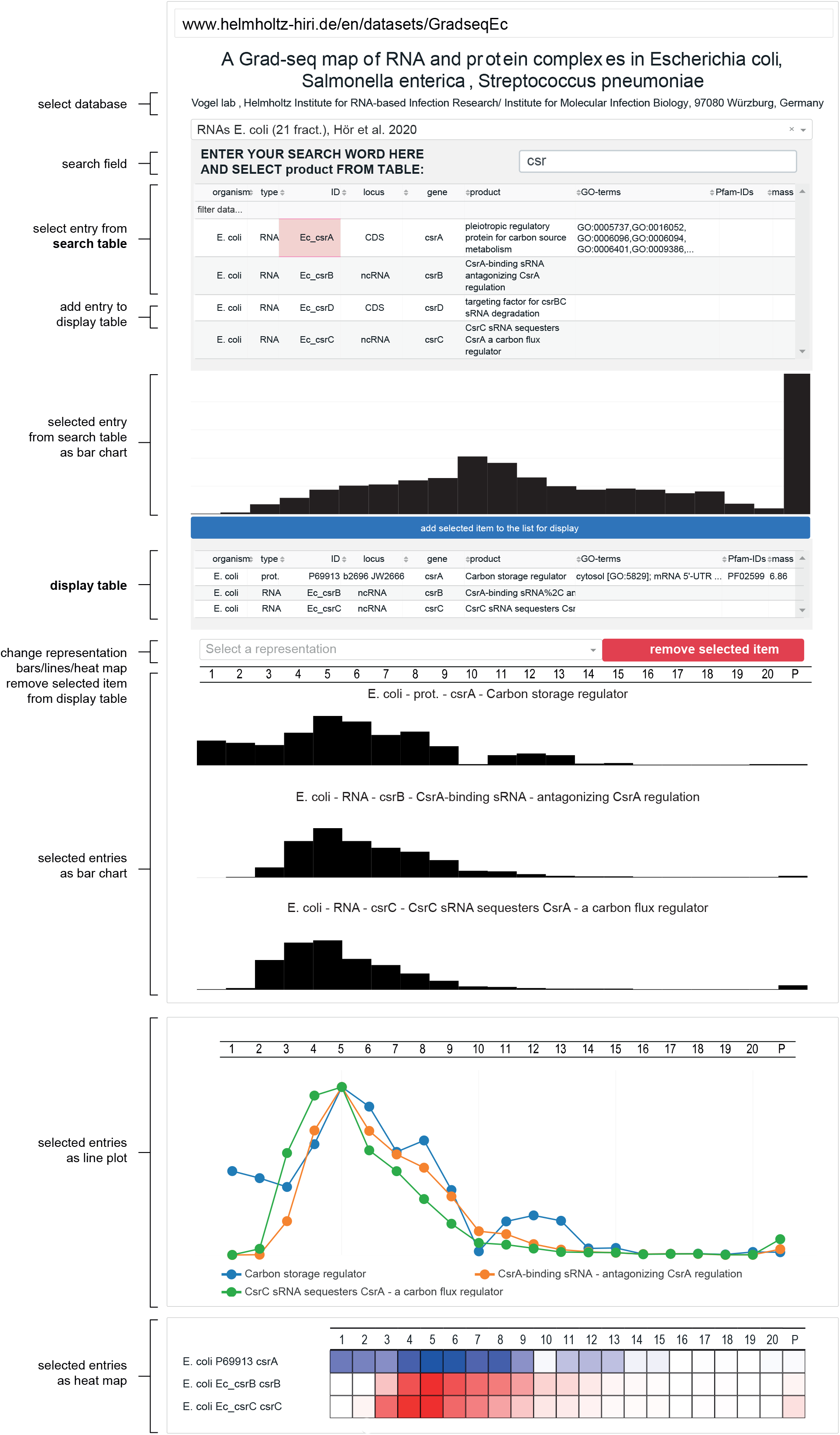
Overview of the Grad-seq online browser, accessible at https://helmholtz-hiri.de/en/datasets/gradseqec/.

## DISCUSSION

Our Grad-seq analysis of *E. coli* provides the first comprehensive landscape of stable RNA and protein complexes in this important model bacterium. As a valuable resource, our Grad-seq dataset adds the previously missing knowledge about potential RNA interactions to the ever-growing pool of information about *E. coli*. Exploring this data, we have identified a phage-encoded, K-12-specific toxic protein as well as a conserved ribosome-binding protein. As more Grad-seq data for other species become available, we will develop a better understanding of the scope of interactions and complexes of RNA and proteins in bacteria.

Although there have been several global studies of the protein interactome of *E. coli* (4–13), global information about RNA complex formation has been lacking. Our Grad-seq data readily reproduce the major cellular RNPs such as the SRP or RNAP, while also providing information about regulatory RBPs such as CsrA, Hfq and ProQ, which together bind the majority of sRNAs within the cell (Figure 1F) (84).

While mRNAs were expected to accumulate in the pellet together with actively translating ribosomes (Supplementary Figure S1A), sRNAs were not (Figure 1F-G and Supplementary Figure S1B-E). Straight-forward explanations for the surprisingly abundant ribosome associations of sRNAs include their activities as activators of mRNA translation (such as DsrA (85,86)) and translational repressors within a polycistronic mRNA (such as Spot 42 (87)). Moreover, ribosome association of sRNAs can hint at a dual function of an RNA, as first describe for *Staphylococcus aureus* RNAIII, which is both a regulatory RNA and the mRNA of δ-hemolysin (88). Here, we observed SgrS—the best-characterized dual-function RNA of *E. coli* (89)—almost exclusively in the pellet fraction (Figure 1F).

The present work highlights strong association with the 30S subunit as a predictor of coding potential. We have discovered that the seemingly noncoding prophage-specific RyeG sRNA encodes a toxic 48 aa protein (Figures 3 and 4), agreeing with recent analysis of ribosome footprints by others (69). At this point, however, we have no indication for a dual function of RyeG, i.e., that the RyeG RNA itself serves as regulator, judging by the fact that a single nucleotide change creating a premature stop codon rendered RyeG non-toxic (Figure 4F). The exceptional 30S subunit association of RyeG could indicate that translation initiation takes place but formation of actively translating ribosomes is somehow inhibited. In our dataset, other mRNAs with potential 30S subunit association (Supplementary Figure S8A) were enriched in pseudogenes, toxins and phage-encoded genes (Supplementary Figure S8B), suggesting there might indeed be a mechanism preventing the translation of non-functional or detrimental RNAs. Alternatively, 30S subunit association could be an intrinsic feature of mRNAs with especially small CDSs that is readily detected by Grad-seq (Supplementary Figure S5C).

The toxicity of RyeG is visible by a much prolonged lag time before bacterial growth takes off after fresh inoculation (Figure 4A and B). Increased lag times can be detrimental to a bacterium because it might be outcompeted by others in the same environment that quicker at utilizing the available nutrients (90). However, lag phase can also be beneficial by conferring stress tolerance to, e.g., antibiotics (91) or by possibly increasing immune evasion (92). The latter might be one benefit of the continuous presence of the defective prophage CPS-53 in the chromosome of *E. coli* K-12, since CPS-53 was shown to increase H_2_O_2_ and acid resistance (66). Yet, the exact function of RyeG and how the cell overcomes its toxic effect remain obscure.

Although our previous Grad-seq analysis of related *Salmonella* bacteria already included the global proteomics component (14), this part remained underexplored. Complementing data obtained via binary co-purification, Grad-seq provides an overview of the major complexes of *E. coli* (Figure 5), and so lends itself to cross-comparison with existing global data obtained from AP/MS and two-hybrid screens (4–8). As a first example, we identified the well-conserved YggL as a 70S ribosome-binding protein (Figure 6A-G), agreeing with binary interactome studies that reported interactions between YggL and the ribosomal proteins L2, S3, S4, L28 and L32 (5,7). Intriguingly, another previous study used a pulse labeling approach in combination with sucrose gradient centrifugation and MS to identify proteins implicated in ribosome assembly, which suggested YggL to be involved in late 50S subunit assembly or final maturation of the 70S ribosome (83). In line with this, we found the knockout of *yggL* to increase the amount of free 50S subunits within the cell (Figure 7H). This implies either an increase in biogenesis of 50S subunits to compensate for a defect in 70S assembly as was observed for other ribosome-associated proteins (93–95).

Our discovery of the ribosome association of YggL further emphasizes the diversity of ribosomes across different species. For example, RbgA is an essential GTPase for 50S assembly in *B. subtilis* (95) but absent from *E. coli*, showing that different species use different proteins for ribosome assembly. Reciprocally, YggL is strongly conserved within the γ-proteobacteria but absent from other bacterial classes. Therefore, might YggL be involved in the formation of subpopulations of specialized ribosomes (96) or exert a function not needed in other groups of bacteria? In this regard, the emerging protein catalogs from Grad-seq in different species promise to yield new types of protein functions in building and shaping the full diversity of bacterial ribosomes.

## DATA AVAILABILITY

The sequencing data have been deposited in NCBI’s Gene Expression Omnibus (97) and are accessible through GEO Series accession number GSE152974 (https://www.ncbi.nlm.nih.gov/geo/query/acc.cgi?acc=GSE152974). The mass spectrometry proteomics data have been deposited to the ProteomeXchange Consortium (98) via the PRIDE (99) partner repository with the dataset identifier PXD019900 (https://www.ebi.ac.uk/pride/archive/projects/PXD019900).

## ACKNOWLEDGMENTS

We thank J.T. Vanselow and A. Schlosser for mass spectrometry analysis, the members of the Vogel lab for splendid ideas and discussions, and V. Herbst for excellent technical assistance.

## SUPPLEMENTARY FIGURE LEGENDS

**Supplementary Figure S1.**
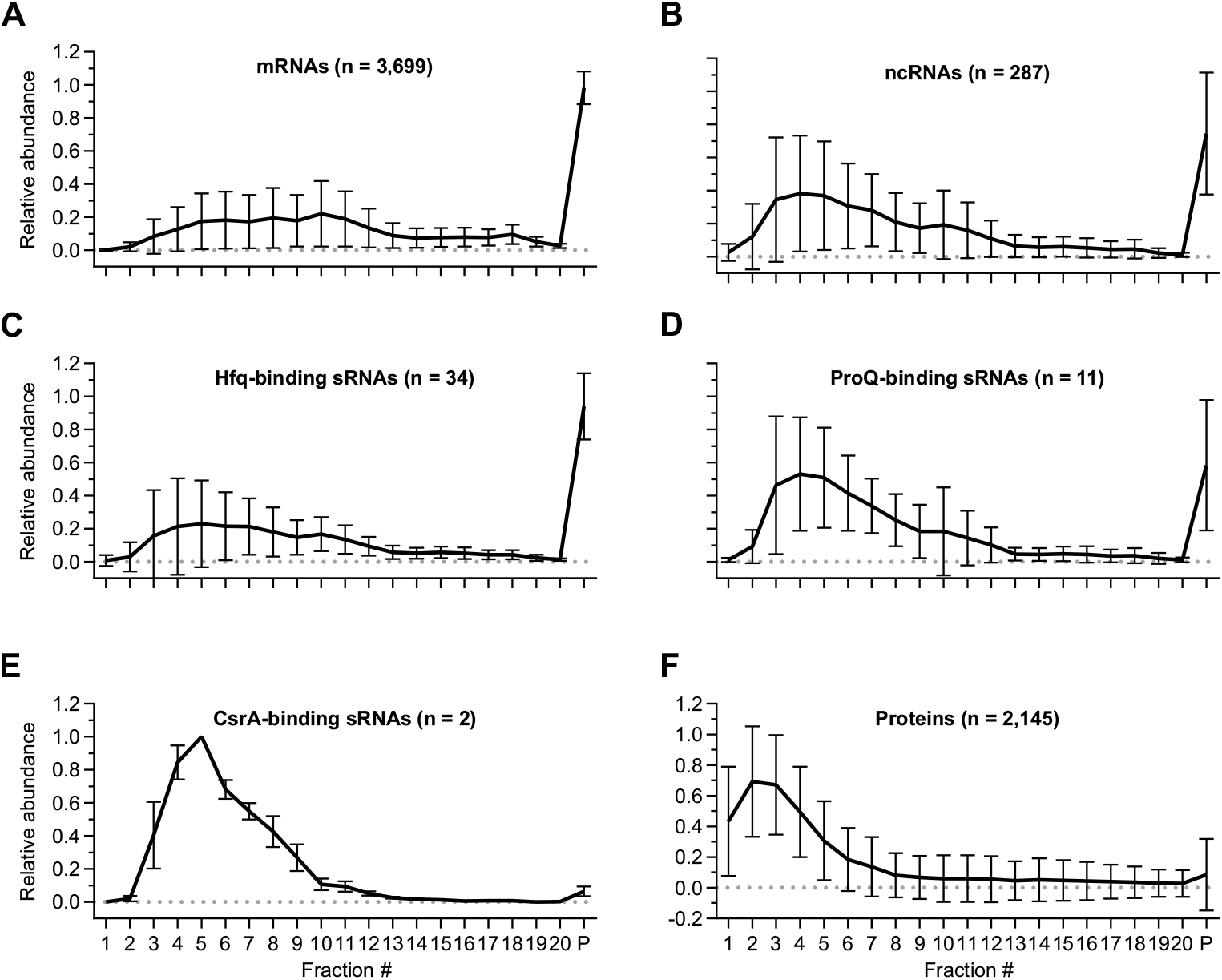
Overview of RNA and protein sedimentation profiles. (A-E) Average sedimentation profiles of all detected mRNAs (A), ncRNAs (B), Hfq-binding sRNAs (C), ProQ-binding sRNAs (D) and CsrA-binding sRNAs (E). sRNAs shown in (C) and (D) were chosen according to the high confidence ones reported in (1). (F) Average sedimentation profiles of all detected proteins. Error bars show SD from the mean.

**Supplementary Figure S2.**
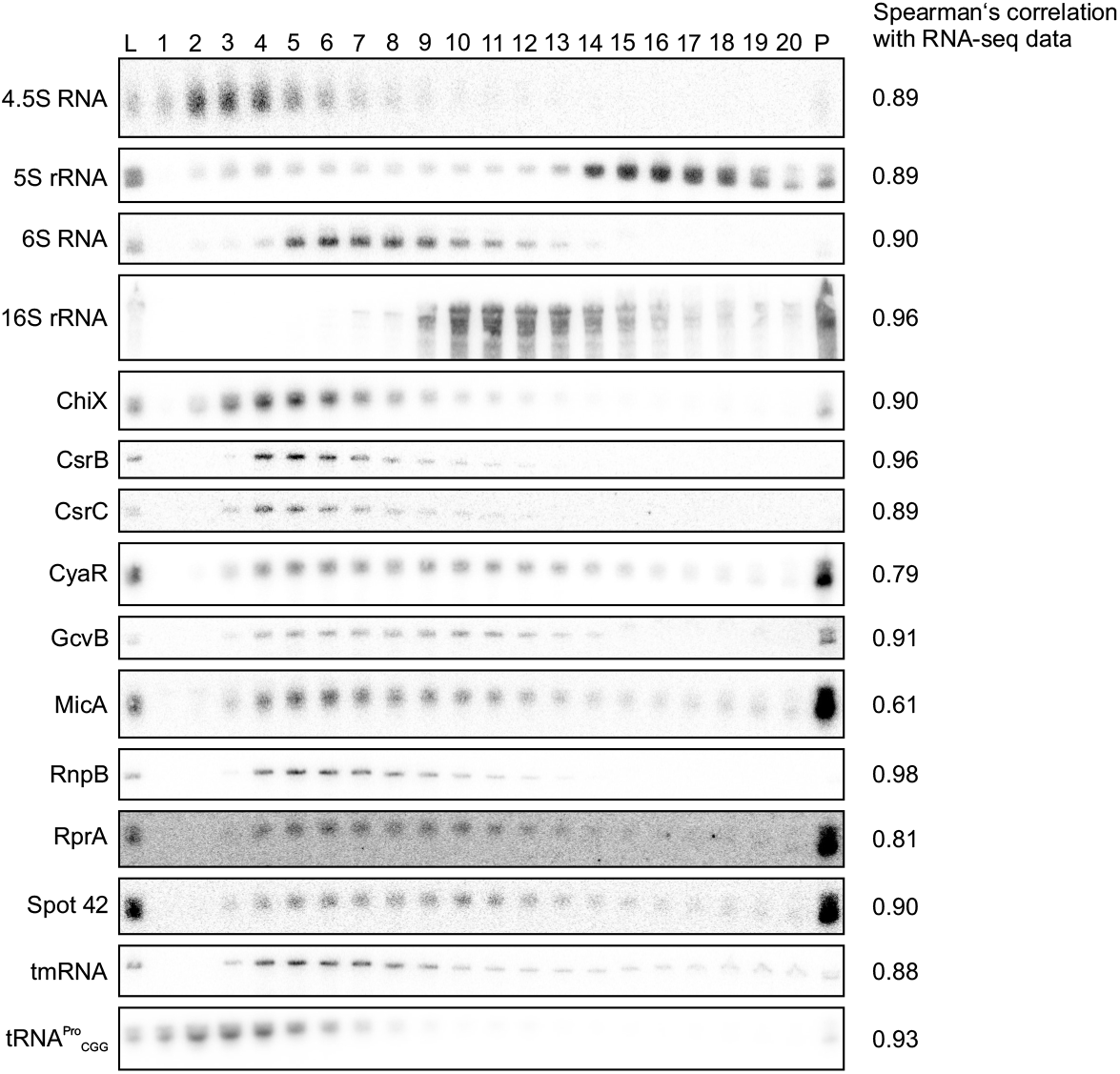
Northern blots of the sedimentation profiles of ncRNAs. Northern blotting reveals different sedimentation profiles for different ncRNAs. Spearman’s correlation of the quantified profiles of each ncRNA with the corresponding RNA-seq data is given on the right.

**Supplementary Figure S3.**
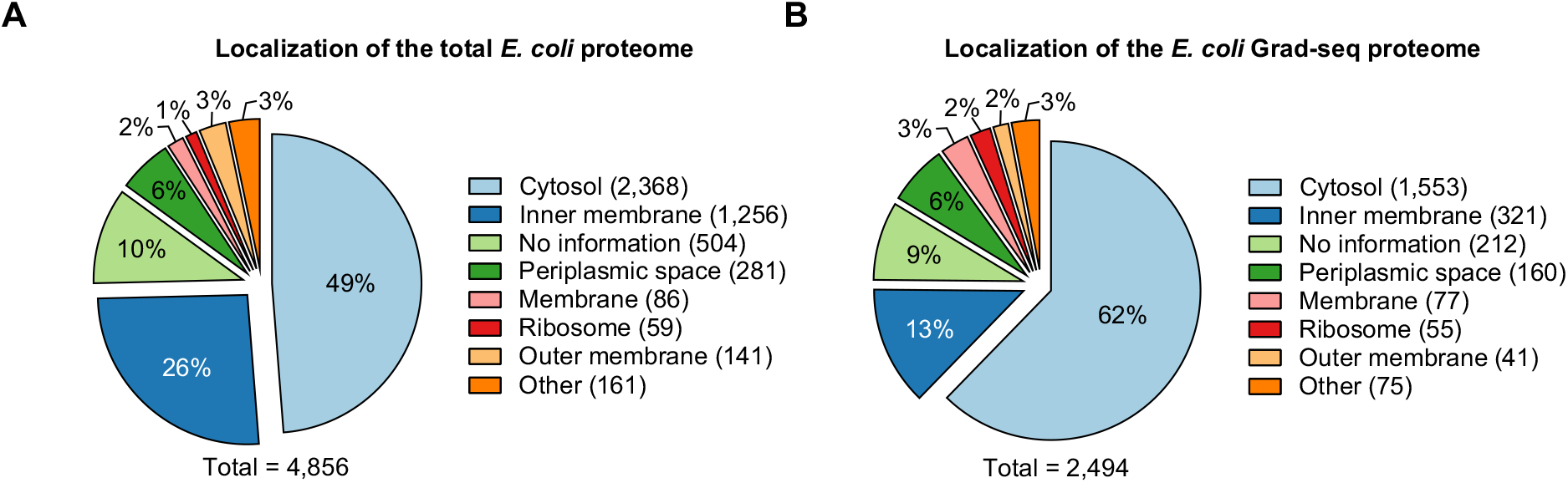
Localization of *E. coli* proteins. (A) Localization of all proteins as reported by EcoCyc (2). (B) Localization of all proteins detected by Grad-seq as reported by EcoCyc (2). Note that the total number of proteins given here does not agree with the number of detected proteins listed in Supplementary Table S2, because some proteins have more than one localization assigned.

**Supplementary Figure S4.**
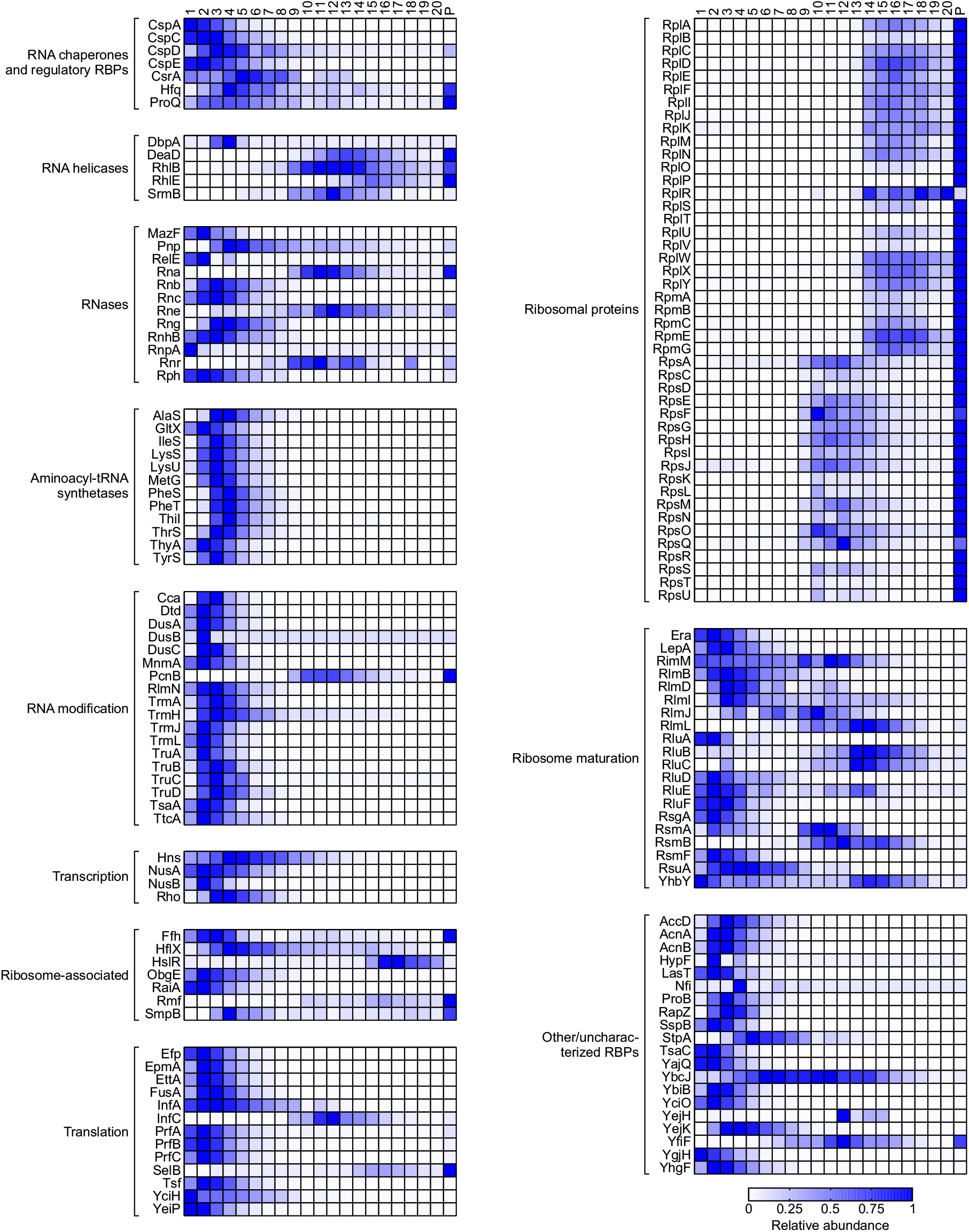
Overview of the sedimentation profiles of RBPs. Heat map showing the sedimentation profiles of 163 proteins with predicted RNA-binding properties based on UniProt (3) and Gene Ontology (4,5) information.

**Supplementary Figure S5.**
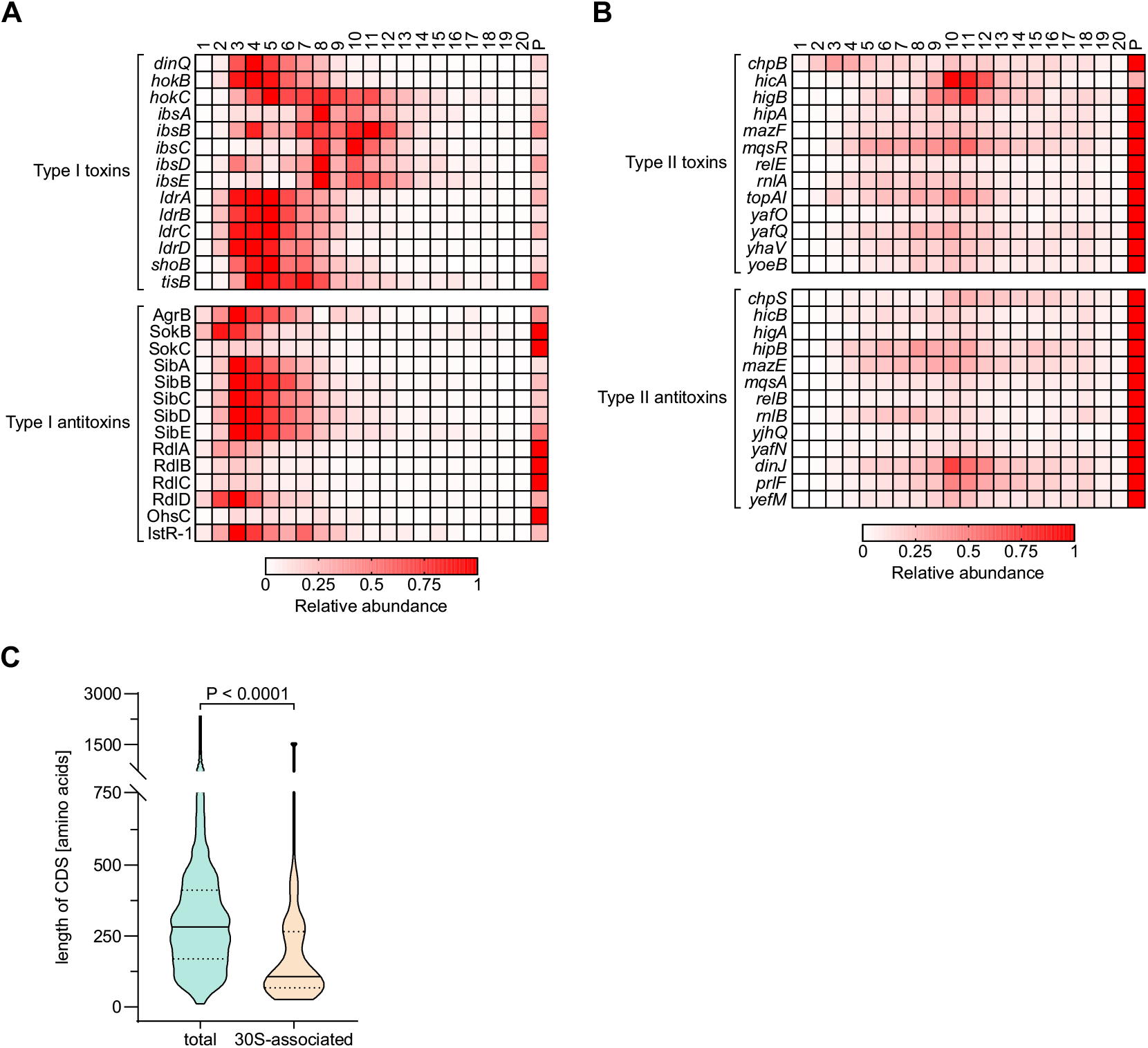
Sedimentation profiles of toxin-antitoxin (TA) system transcripts. (A) Heat map showing the sedimentation profiles of type I TA system toxin mRNAs (top) and their corresponding antitoxin ncRNAs (bottom). (B) Heat map showing the sedimentation profiles of type II TA system toxin mRNAs (top) and their corresponding antitoxin mRNAs (bottom). (C) Violin plot showing the length distribution of all non-pseudo CDSs of *E. coli* (green, n = 4,141) and the non-pseudo CDSs of the 30S-associated mRNAs shown in Figure 2B (orange, n = 29). The solid line indicates the median, whereas the dashed lines indicate the upper and lower quartiles. P-value was calculated using a Kolmogorov-Smirnov test.

**Supplementary Figure S6.**
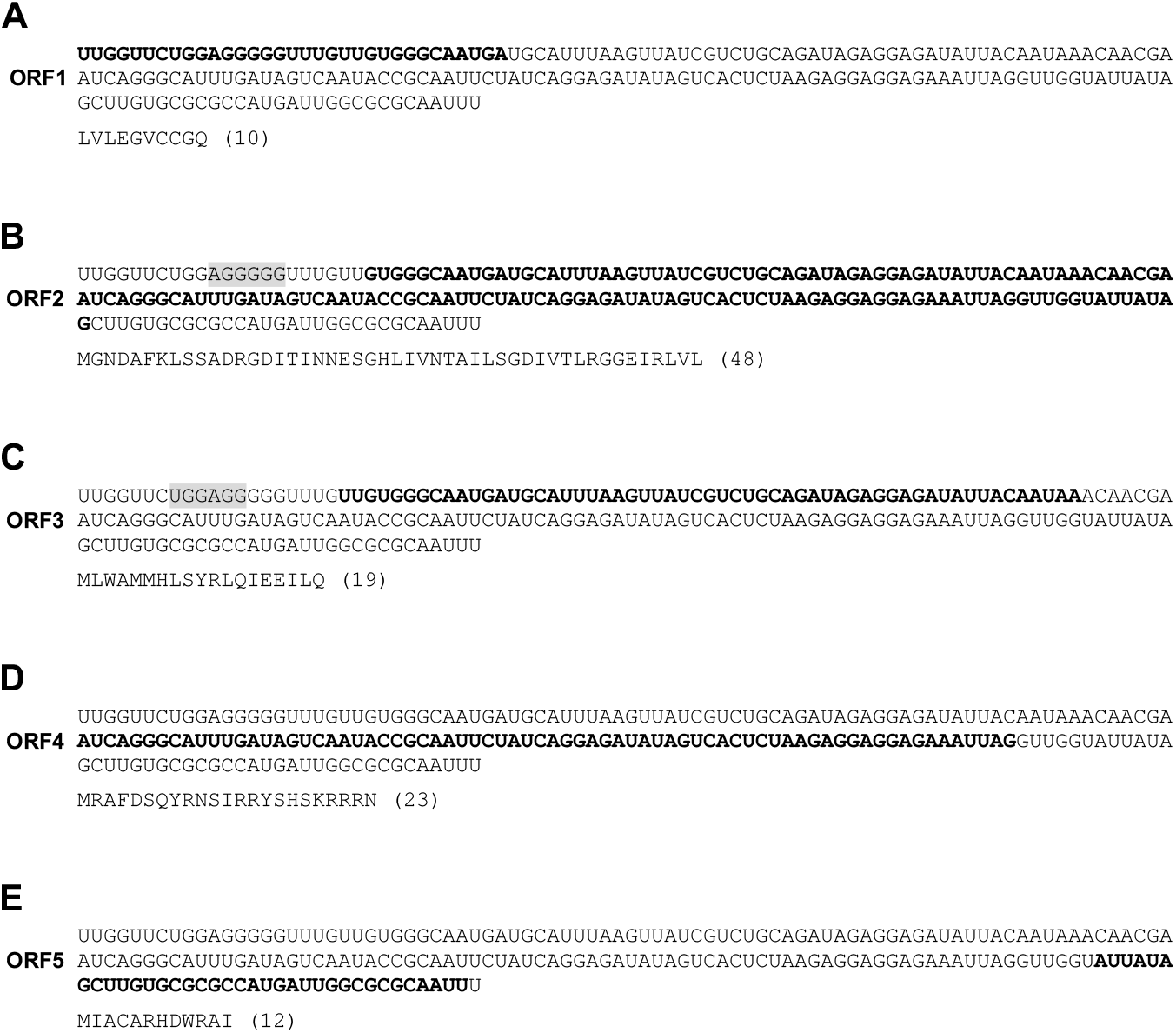
Predicted ORFs within RyeG. (A-E) All five ORFs of RyeG that were predicted by ORFfinder (https://www.ncbi.nlm.nih.gov/orffinder/) are highlighted in bold. The corresponding amino acid sequences are given below each corresponding ORF. For ORF2 (B) and ORF3 (C), predicted ribosome-binding sites are highlighted in gray.

**Supplementary Figure S7.**
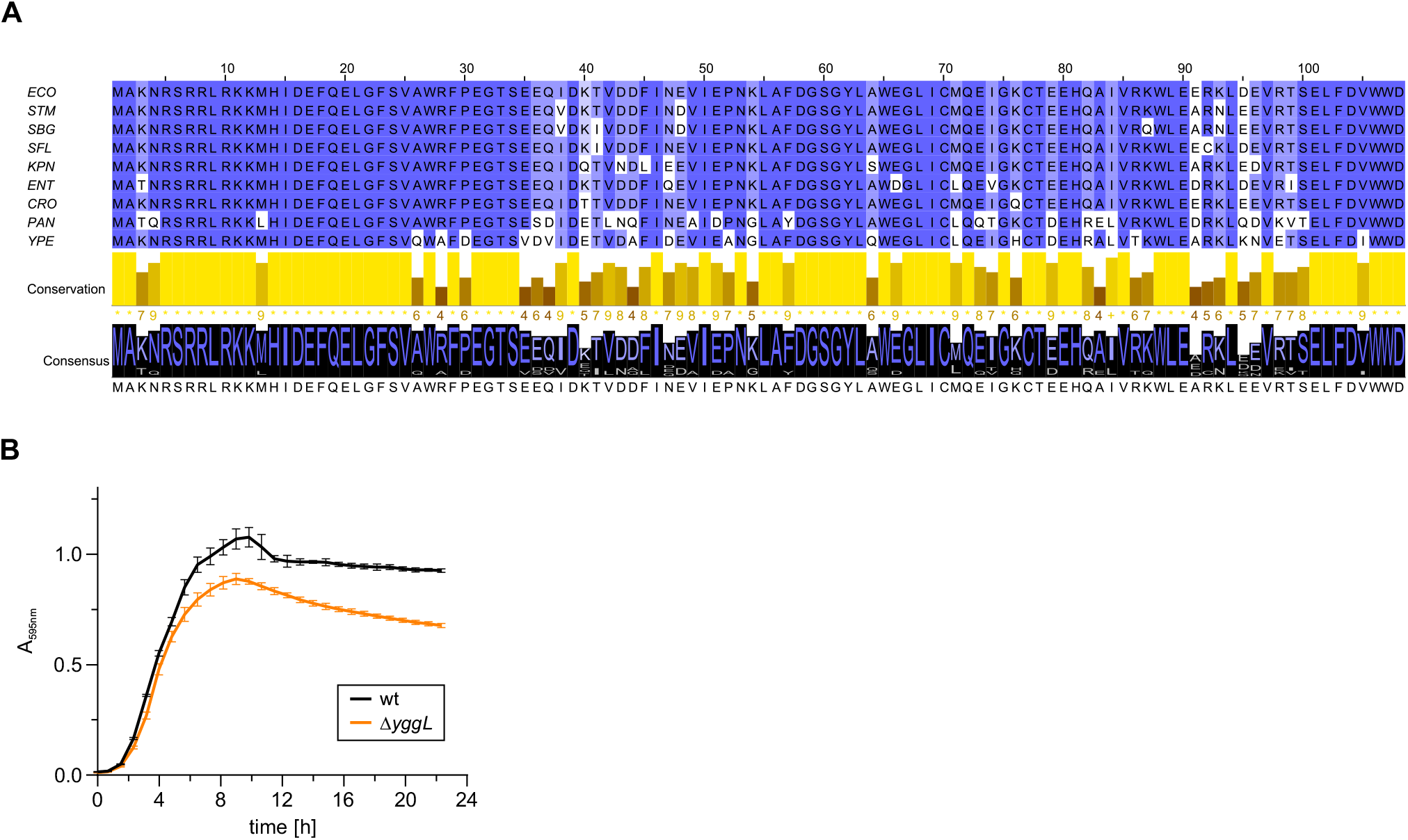
YggL is a conserved protein that binds the 50S subunit of the 70S ribosome. (A) Multiple sequence alignment of YggL homologs from *E. coli* (ECO), *Salmonella* Typhimurium (STM), *Salmonella bongori* (SBG), *Shigella flexneri* (SFL), *Klebsiella pneumoniae* (KPN), *Enterobacter cloacae* (ENT), *Citrobacter rodentium* (CRO), *Pantoea* spp. (PAN) and *Yersinia pestis* (YPE) using Clustal Omega (6). YggL is highly conserved on amino acid level between different members of γ-proteobacteria. Residues with ≥ 50% identity are highlighted in a blue gradient. Visualization was performed using Jalview (7). (B) Growth curves comparing wild-type *E. coli* to Δ*yggL*. No difference in growth can be detected between the strains. Data obtained from three biological replicates. Error bars show SD from the mean.

**Supplementary Figure S8.**
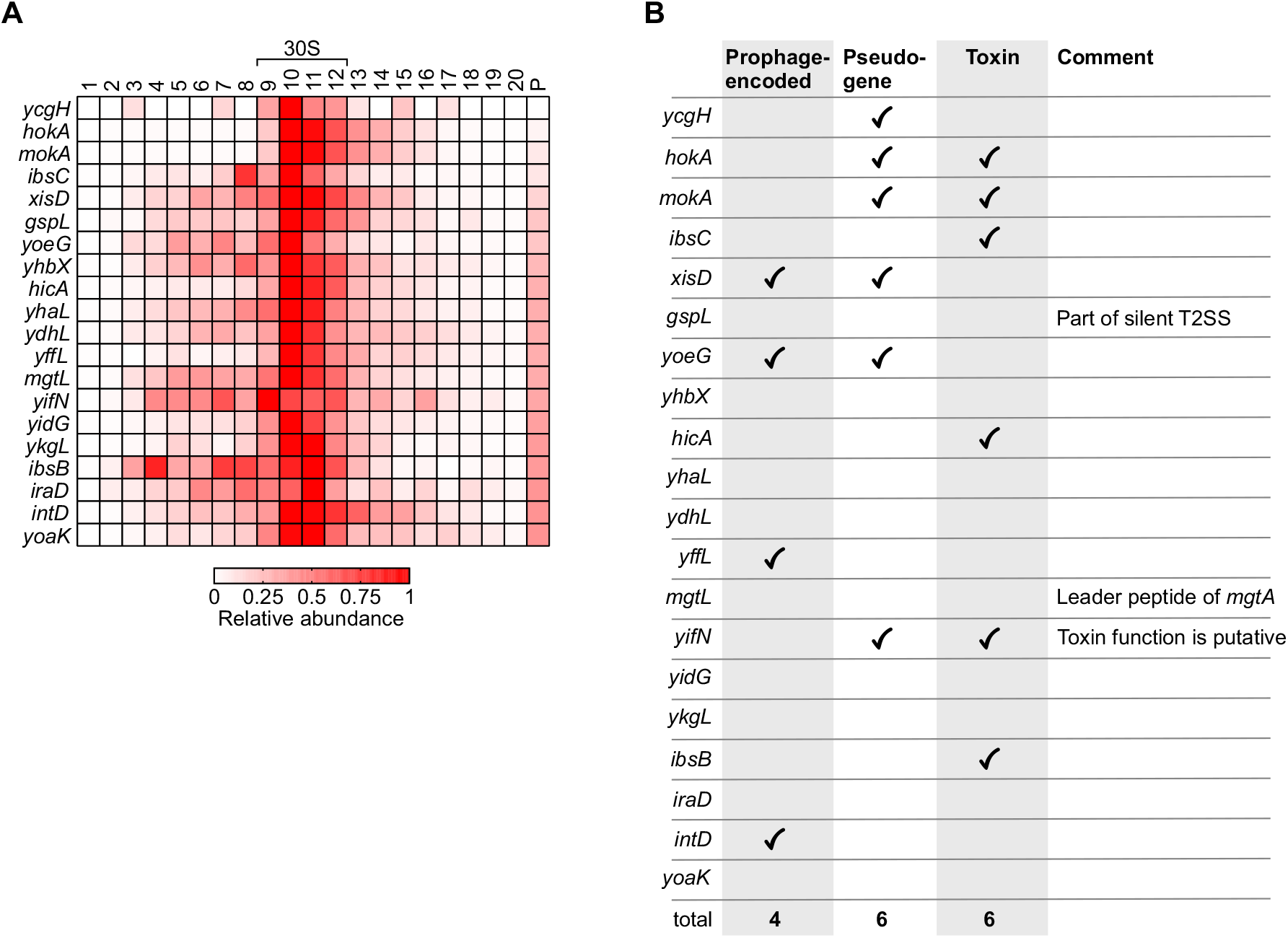
30S subunit-associated mRNAs are enriched in horizontally acquired, non-functional and toxic CDSs. (A) Heat map showing the top 20 mRNAs found to have their peak abundance at the 30S subunit (fractions 9-12), sorted by increasing pellet abundance. (B) List of the 20 mRNAs shown in (A) with added information about them being prophage-encoded, a pseudogene or a toxin (obtained from EcoCyc (2)). T2SS, type 2 secretion system.

